# Targeting H3K4 methylation as a novel therapeutic strategy against tumor infiltration and nuclear changes of acute lymphoblastic leukemia cells

**DOI:** 10.1101/2022.06.16.495903

**Authors:** Raquel González-Novo, Ana de Lope-Planelles, África González-Murillo, Elena Madrazo, David Acitores, Mario García de Lacoba, Manuel Ramírez, Javier Redondo-Muñoz

## Abstract

Acute lymphoblastic leukemia (ALL) is the most common pediatric cancer, and the infiltration of leukemic cells is critical for disease progression and relapse. In spite of the canonical functions of histone methylation in gene regulation, differentiation, and DNA homeostasis; its contribution to the nuclear deformability of migrating leukemic cells remains unclear. Here, we showed that 3D conditions promoted a fast upregulation of H3K4 methylation, bound to transcriptional changes in ALL cells. Furthermore, we demonstrated that targeting WDR5 (a core subunit involved in H3K4 methylation) impaired the invasion of leukemia cells in vitro, and their tissue infiltration in an immunodeficient mouse model. WDR5 expression correlated with other cell receptors involved in leukemia dissemination in clinical samples from ALL patients. Interestingly, blocking WDR5 did not reduce the chemotactic response of leukemia cells, suggesting a different mechanism by which H3K4 methylation might operate at both nuclear and functional level to control ALL cell invasiveness in 3D conditions. We applied biochemical and biophysical approaches to determine that H3K4 methylation induced by 3D conditions was dependent on MLCK activity, and regulated the chromatin compaction and the mechanical nuclear response of leukemia cells in 3D conditions. Collectively, our data revealed that confined conditions provide novel molecular and biophysical mechanisms used by leukemia cells to disseminate, suggesting H3K4 methylation and nuclear mechanical pathways as promising therapeutic targets against ALL infiltration.

**Highlights:** 3D conditions induce H3K4 methylation and transcriptional changes in ALL cells.

Targeting WDR5 and H3K4 methylation blocks ALL cell invasion in vitro 3D conditions and leukemia dissemination in vivo.

WDR5 expression correlates with other cell receptors related to leukemia migration in clinical samples from patients with ALL.

H3K4 methylation induced by 3D conditions is dependent of MLCK activity and regulates cell movement through 3D environments.

Leukemia cells in 3D conditions alter their chromatin compaction and the biomechanical deformability of their nuclei.

## Introduction

Acute lymphoblastic leukemia (ALL) is the most common pediatric cancer; despite current therapies having improved the outcomes for children with ALL up to 90%, a proportion of patients still experience therapies failure and fatal relapse.^1^ Leukemic cells interact with their surrounding environment, which provides biochemical and physical signals that control leukemia initiation, proliferation, survival, and tissue infiltration.^2^ Migration through these 3D environments promotes nuclear alterations such as gene regulation, epigenetic changes and DNA damage markers.^3,4^ This is particularly important for the nucleus, which acts as a cellular mechanosensor to allow cancer cells to deform and to squeeze through physical barriers, such as the interstitial space and endothelial barriers during their infiltration into other organs.^5,6^ Genetic and epigenetic studies have identified potential prognosis markers and therapeutic targets against leukemia initiation and progression.^7,8^ However, the interplay between the surrounding environment and epigenetic changes induced in ALL cells remain functionally unexplored.

Histone methylation plays a critical role in chromatin compaction, gene regulation, DNA recombination and repair of cancer cells.^9^ The MLL (mixed-lineage leukemia) methyltransferases associate with the WRAD complex (comprising WDR5, RBP5, ASH2L, and DPY-30) to catalyze the histone H3 lysine 4 (H3K4) methylation.^10^ Recent works have demonstrated that WDR5 is involved in the malignant transformation and progression of multiple cancers, including leukemia.^11,12^ In addition, WDR5 regulates breast cancer metastasis and lymphocyte migration.^13,14^

Understanding whether epigenetic changes are required for ALL progression and dissemination would foster therapeutic strategies targeting their functional effects. To directly address this, we identified how external 3D conditions promoted upregulation of H3K4 methylation in ALL cells, concomitant with epigenomic and transcriptional changes in genes involved in several leukemia-related progression pathways. We demonstrated that targeting WDR5 reduced leukemia cell infiltration and migration *in vitro* and *in vivo*. Notably, although we found a correlation between the expression of WDR5 and other cell receptors involved in ALL migration, WDR5 inhibition did not affect the chemotactic response of ALL cells, suggesting a fundamental role for the cell migration in 3D conditions. We performed a biophysical and molecular characterization of the global chromatin structure and determined that 3D conditions induced global changes in the nucleus of ALL cells. Furthermore, we defined that ALL cells in suspension or embedded in 3D presented a different biophysical signature of their nuclei, which is critical for nuclear deformability and migration across confined spaces. Together, our results strengthen new insights regarding a potential therapeutic target of H3K4 methylation in the biomechanical response and invasion of leukemia cells.

## Materials and Methods

### Cells and cell culture

The human CCRF-CEM and Reh ALL cell lines were obtained from Dr. Ramírez (Hospital Infantil Universitario Niño Jesús). ALL primary cells were obtained from patients under 16 years old with informed consent for research purposes at Hospital Universitario Niño Jesús (**Supplementary Table 1**). ALL diagnosis and treatment were defined according to SEHOP-PETHEMA 2013 (Spanish Program for the Treatment of Hematologic Diseases). All cells were cultured in RPMI 1640 medium with L-glutamine and 25 mM Hepes (Sigma Aldrich, St. Louis, MO, USA) and 10% fetal bovine serum (Sigma-Aldrich) and maintained in 5% CO_2_ and 37°C.

Cells were cultured in suspension, on plates coated with VCAM-1 or embedded in a collagen matrix. For 3D matrix, collagen type I from bovine were reconstituted at 1.7 mg/mL in RPMI and neutralized with 7.5% NaHCO3 and 25 mM Hepes. Cells were added to the collagen solution before polymerization at 37 °C for 1 h. In some cases, cells were preincubated with specific chemical inhibitors and the medium was also supplemented with these inhibitors.

### Cell penetration assay

A 100 µl collagen matrix was reconstituted at 1.7 mg/mL in RPMI, neutralized with 7.5% NaHCO3 and 25 mM Hepes and allowed to polymerize inside transwell inserts (5 µm, Costar). 3 × 10^5^ cells were pretreated or not with OICR-9429 for 1 h in serum-free RPMI, or transfected with control-and WDR5-shRNAs for 24 h before seeding onto a collagen gel. RPMI medium with 10% of FBS was added to the bottom chamber of the transwell as chemoattractant. After 24 h, invading cells were fixed with 4% PFA for 1 h, permeabilized with 0.5% Triton-X-100 in PBS for 30 min and stained with propidium iodide. Invading cells were imaged with a sCMOS Orca-Flash 4.0LT camera (Hamamatsu) coupled 5 to an inverted DMi8 microscope (Leica), capturing serial z-stacks every 10 µm with a 10× objective (dry ACS APO 10x/NA 0.3). Percentage of penetrating cells in each range of distances was quantified with FIJI software (National Institute Health, US).

### In-vivo short-term leukemia homing assay

NOD-SCID-Il2rg-/-(NSG) mice (*Mus musculus*), were purchased at Charles River (France), and bred and maintained at the Servicio del Animalario del Centro de Investigaciones Energéticas, Medioambientales y Tecnológicas (CIEMAT) with number 28079-21 A. All mice were used following guidelines issued by the European and Spanish legislations for laboratory animal care. 5× 10^6^ control and WDR5 inhibited CCRF-CEM cells were labeled with Cell Tracker Far Red (1 µM) and CFSE (5 µM), respectively. After 30 min, cells were mixed and intravenously (IV) administered to 15 weeks-old non-conditioned NSG mice. Sacrifice was performed 3 hours after injection. Bone marrow from femurs and spleens were extracted, processed through mechanical disaggregation and stained cells were resuspended in the PBS. Samples were acquired in a FacsCanto II (BD, San Jose, CA) cytometer and the number of labeled cells analyzed using FacsDiva software. Transfected control (labeled with Cell Tracker Far Red) and WDR5 shRNA (labeled with CFSE) CCRF-CEM cells were mixed and intravenously administered to NSG mice. Organs were extracted, processed through mechanical disaggregation and stained cells were resuspended in the PBS and analyzed by flow cytometry. Also, the distribution among the tissues of colonizing cells was studied by sample fixation in formaldehyde, frozen in OCT (optimum cutting temperature) and serial cryostat sections were stained with Dapi and directly imaged with a 20× objective.

## Results

### The 3D environment induces changes in the genome-wide H3K4 methylation and transcriptional program of leukemia cells

We first cultured ALL cell lines in suspension and embedded them in a 3D collagen matrix to determine the contribution of the microenvironment to the H3K4 methylation in leukemia cells. In both cell types, the 3D environment upregulated the levels of H3K4me3 (**Figures 1A, 1B**). We confirmed that 3D culture conditions did not alter the expression of other histone markers and c-Myc in ALL cells at the times studied (**Figure S1A**). To determine the changes in the levels and pattern of H3K4 methylation on chromosomes, we performed H3K4me3 ChIP-seq (chromatin immunoprecipitation sequencing) analysis in CCRF-CEM cells cultured in suspension (control) and 3D conditions and found that the predominant categories enriched in H3K4me3 peaks corresponded to those genes related with cell differentiation and cycle, intracellular signaling, oxidative stress and metabolism (**Figure 1C**). Since H3K4 methylation is linked to active gene transcription,^15^ we hypothesized that cell confinement may promote transcriptional changes related to leukemia progression. To address this, we performed expression profiling on a set of RNAs from CCRF-CEM cells cultured in suspension or embedded in 3D conditions. We identified the differential expression of 163 up-and 84 down-regulated genes in CCRF-CEM cells in 3D conditions with a cut-off of |Fold Change| > 1.4 and a P-value of 0.05 from 21448 genes (**Figure 1D, Table S1**). Our Gene Ontology (GO) enrichment analysis showed changes in transcripts related to transcriptional regulation, cell division and mitosis and DNA repair (**Figure S1B**). To determine the possible association between the global change of H3K4 methylation induced by 3D culture conditions and its potential transcriptional role, we compared the features of the microarray and the ChIPseq analysis (**Figure S1C**). Interestingly, we did not find a correlation between H3K4me3 occupancy and the transcriptional changes, suggesting that the H3K4me3 upregulation induced by 3D conditions might be regulating additional cellular functions (**Figure 1E**). Together, we conclude that 3D culture conditions promoted an upregulation of H3K4me3 levels and transcriptional changes in leukemia cells.

**Fig. 1.**
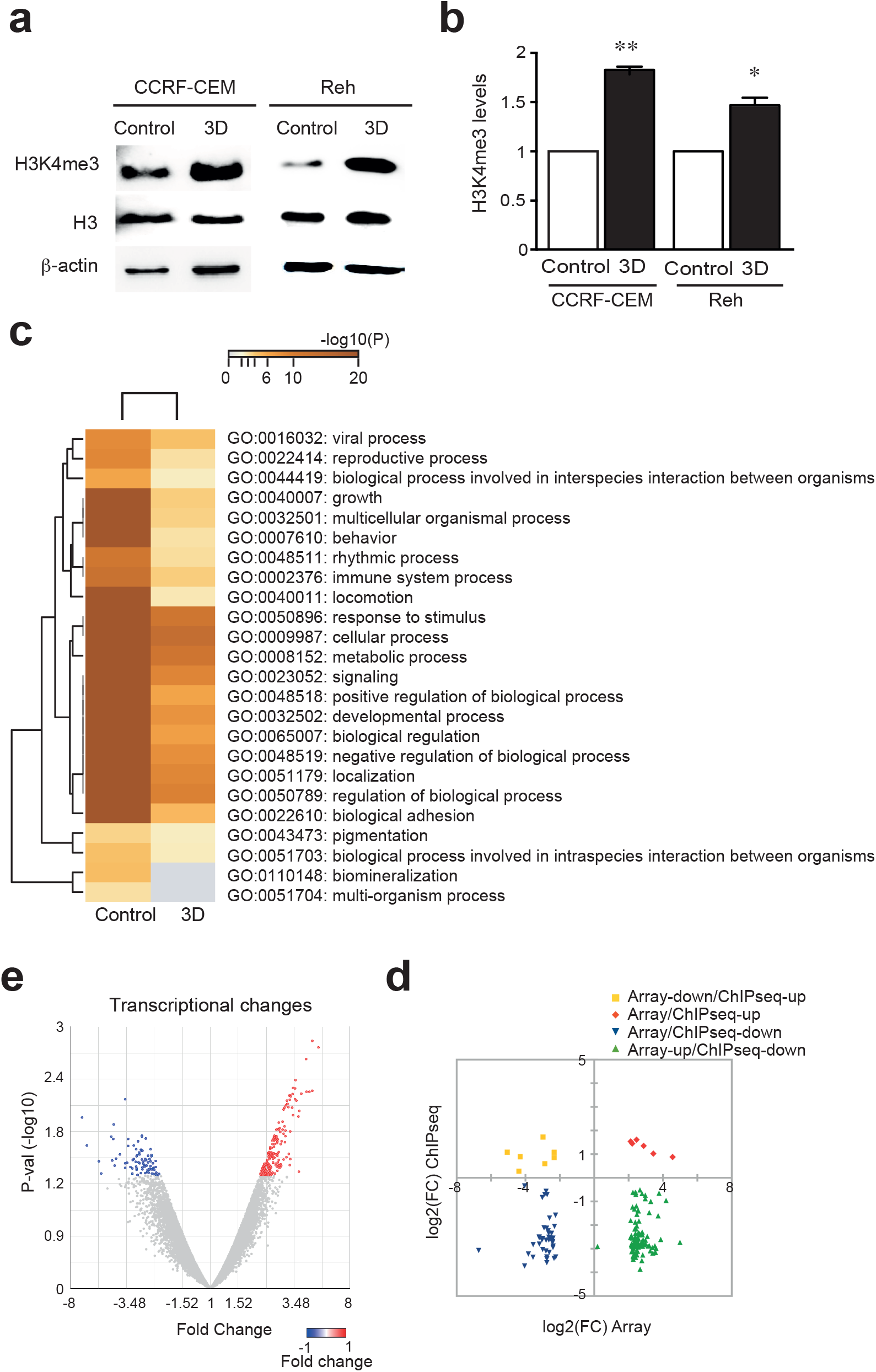
3D conditions induced changes in the genome-wide H3K4 methylation of leukemia cells. **(A)** CCRF-CEM and Reh cells were cultured in suspension (Control) or embedded in a 3D collagen matrix (3D). After 1 h, cells were collected and H3K4me3 levels tested by western blotting. **(B)** Graph shows the H3K4 methylation levels normalized to loading controls. Mean n = 3 ± SEM. **(C)** CCRF-CEM cells were cultured in suspension (control) or in a 3D collagen matrix (3D) for 1h. Then, cells were collected and H3K4me3 ChIPseq assay was performed. Graph shows the top gene ontology enrichment results of H3K4me3 ChIPseq assay. **(D)** CCRF-CEM cells were cultured in suspension or 3D conditions. RNA was isolated and the number of differentially expressed genes was analyzed by microarray. Volcano plots show significantly upregulated transcripts in control (blue) and 3D (red) conditions. **(E)** Correlation plots of differentially expressed transcripts and H3K4me3 occupancy from ChIPseq assays in Jurkat and CCRF-CEM cultured in 3D conditions. (*) P < 0.05, (**) P < 0.01.

### Blocking H3K4 methylation impaired the penetration of ALL cells in 3D confined conditions

To determine the functional consequences of H3K4 methylation on leukemia invasiveness, we cultured ALL cells onto the surface of 3D collagen matrices to analyze their infiltration upon treatment with a specific inhibitor against H3K4 methylation and the WDR5-MLL interaction, called OICR-9429.^16^ We visualized by confocal sections the invasiveness of ALL cells (**Figures 2A, S2A**) and found that OICR-9429 treatment impaired the cell penetration in primary ALL samples (**Figures 2B and 2C**) and both ALL cell lines (**Figures S2B and S2C**). To discard that OICR-9429 treatment might be affecting the cell migration via cell cycle regulation or apoptosis, we cultured ALL cell lines in the presence of OICR-9429 and confirmed that the cell cycle progression and survival of CCRF-CEM and Reh cells were not affected at 3h (data not shown) and 24 h (**Figures 2D and 2E**) upon OICR-9429 treatment. Accordingly, we did not find significant differences in the expression of cyclins or p-H2AX upon OICR-9429 treatment (**Figure 2F**). Taken together, these data suggested that WDR5 and its H3K4 methylation activity were required for ALL cell infiltration and migration into a 3D collagen matrix.

**Fig. 2.**
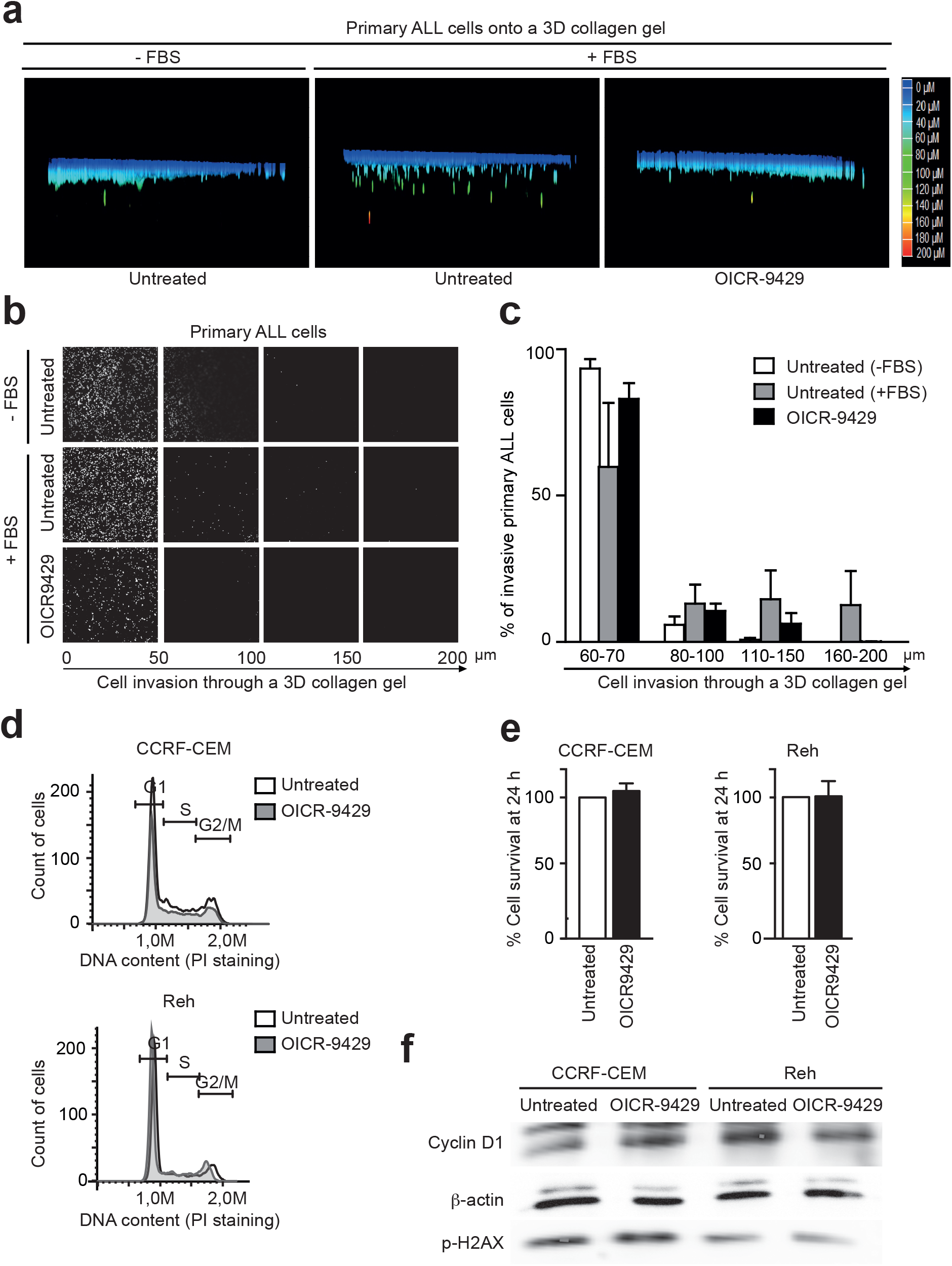
Targeting H3K4 methylation reduces the leukemia invasiveness in 3D conditions. **(A)** ALL primary cells were pretreated or not with 1 μM OICR-9429 (WDR5-MLL inhibitor) for 1h, then cells were seeded on the top of collagen matrix and allowed to penetrate into the collagen in response to serum (FBS, fetal bovine serum) for 24h. Cells were fixed, stained with Propidium Iodide and serial confocal sections were captured. Images show the cell penetrability into the 3D collagen matrix. **(B)** Representative serial confocal sections were captured from cells stained as in (A). **(C)** Graphs show the percentage of cell invasion into the collagen gel in response to serum. Mean n = 3 ± SEM. **(D)** CCRF-CEM and Reh cells were treated with 1 μM OICR-9429 for 24h and then the cell cycle was analyzed by flow cytometry. Graphs show the G1, S and G2/M phases according to DNA content. **(E)** CCRF-CEM and Reh cells were treated with 1 μM OICR-9429 for 24h. Then, cells were collected, fixed and cell viability was determined by Annexin V and PI staining. **(F)** CCRF-CEM and Reh cells were cultured in the presence of OICR-9429 for 24 h, lysed and the levels of cyclin D1 (G1 phase marker) and p-H2AX (DNA damage marker) were resolved by western blotting.

### WDR5 inhibition impairs the infiltration of ALL cells *in vivo*

To enhance the physiological relevance of targeting WDR5 during ALL migration, we mixed control and OICR-9429-treated cells and injected them into the tail vein of recipient immunodeficient NSG (NOD scid gamma) mice. Mice were sacrificed after 3 h and the presence of ALL cells in the bone marrow and the spleen were assessed by organ disaggregation and flow cytometry (**Figure 3A**). We observed that OICR-9429 pretreatment impaired the homing of ALL cells into the spleen (**Figure 3B**). As OICR-9429 treatment impaired the cell migration in vitro and in vivo, we depleted WDR5 expression in both cell lines (**Figure 3C**). We confirmed that WDR5 depletion reduced the number of penetrating ALL cells into the collagen matrix (**Figure S3A**). Next, we further investigated whether WDR5 depletion might affect the cell cycle and DNA damage markers at the studied times, and found no significant changes in the levels of cyclin D1 or p-H2AX upon WDR5 silencing, nor the cell cycle progression (**Figures S3B S3C**). However, WDR5 silencing showed a more remarkable reduction than OICR-9429 treatment in the homing of ALL cells into the spleen and the bone marrow (**Figure 3D)**. When we analyzed the specific infiltration and localization of ALL cells within different organs, we did not observe significant differences in the distribution throughout the tissues of infiltrated ALL cells (**Figures 3E and 3F**). Together, our data suggest that WDR5 activity is fundamental during leukemia cell dissemination within the organism.

**Fig. 3.**
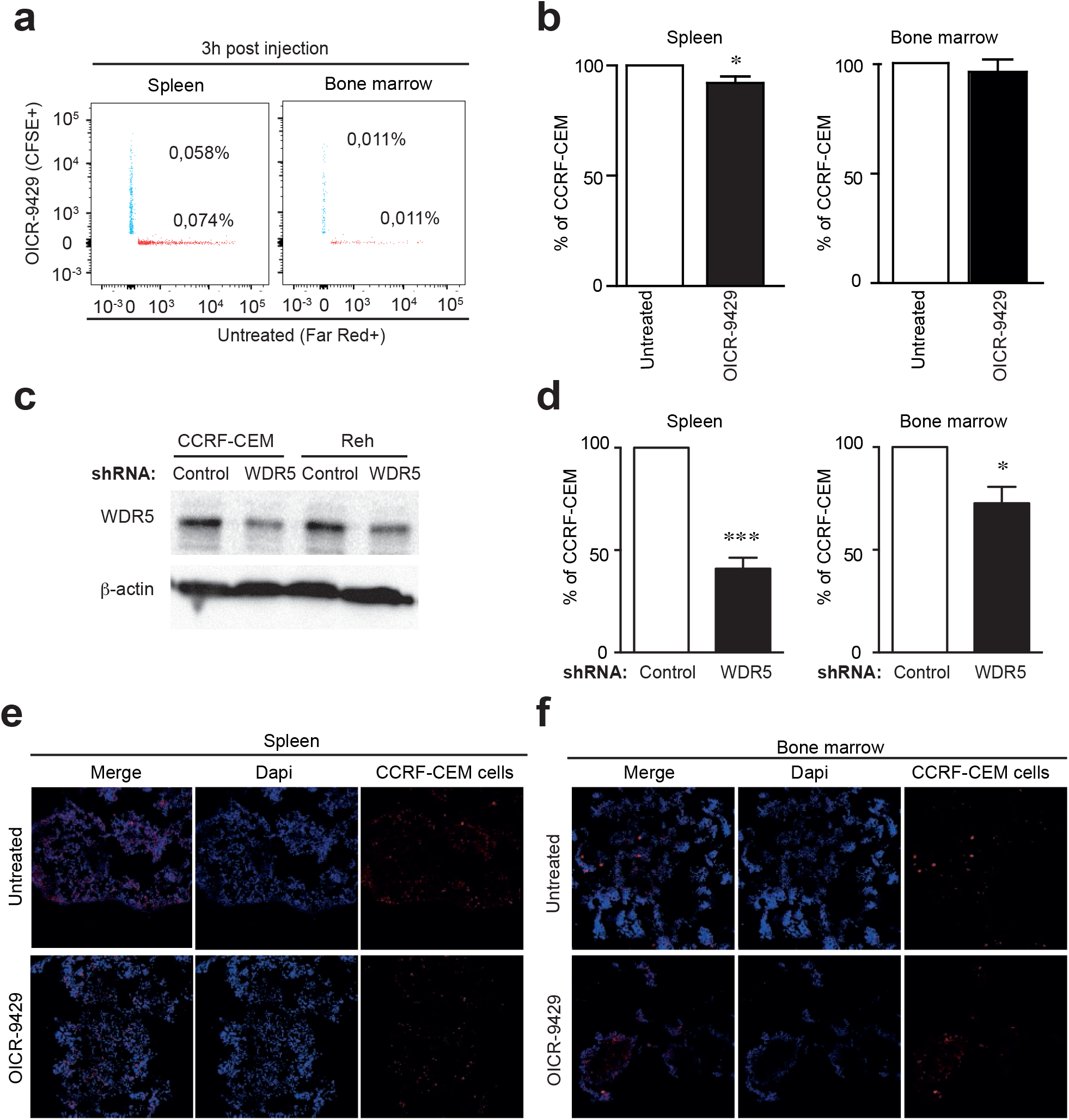
WDR5 controls tissue infiltration of ALL cells in an *in vivo* mouse system. **(A)** CCRF-CEM cells pretreated or not with 1 µM OICR-9429 were labeled with 1 mM Cell Tracker Far Red (control) or 5 mM CFSE (OICR-9429 treated) and injected into the tail vein of NSG mice (n=4). After 3 h, labeled cells in spleen and bone marrow were quantified by flow cytometry. Numbers indicate the percentage of labeled cells according to the total number of events. **(B)** Graph shows the percentage of cells in organs normalized to control (untreated) cells. Mean n = 4 ± SEM. (*) P < 0.05, (**) P < 0.01. **(C)** CCRF-CEM and Reh cells were transfected with control or WDR5 shRNAs for 24 h. Then, cells were collected and WDR5 knocking-down was confirmed by western blotting. (**D**) CCRF-CEM cells transfected with WDR5 shRNA (Cell Tracker Far Red^+^) or with control shRNA (CFSE^+^) were mixed and injected into the tail vein of NSG mice (n=4). After 3 h, labeled cells in spleen and bone marrow were determined by flow cytometry. Graph shows the percentage of labeled cells according to the total number of events. Mean n = 4 ± SEM. **(E, F)** Representative images of confocal sections of spleen (E) and bone marrow (F) from mice injected with CCRF-CEM as in (a). (*) P < 0.05, (**) P < 0.01, (***) P < 0.001.

### WDR5 expression is associated with cell migration molecules in samples of ALL patients

Leukemia cell migration depends on integrins and chemokine receptors,^3^ and impaired ALL infiltration might be related to defects in how ALL cells respond to chemotactic gradients. Interestingly, OICR-9429 treatment of primary samples and ALL cell lines did not reduce the migration towards chemoattractant (**Figure 4A)**. We also confirmed that WDR5 depletion did not reduce the chemotactic migration of CCRF-CEM and Reh cells (**Figure S4A**). To determine how WDR5 expression might be linked to clinical features and adhesive molecules, we analyzed the expression of WDR5 in primary samples from ALL patients (**Table 1)**. We found that WDR5 levels were not related to age or gender of patients with ALL (data not shown), although we observed significant differences in those patients with more than 80% of blast in the bone marrow at diagnosis (**Figure S4B**). We found a positive correlation between the mRNA levels of WDR5 and the receptor CXCR4 (C-X-C Motif Chemokine Receptor 4) in primary samples from patients with ALL (**Figure 4B**). When we compared WDR5 expression and the mRNA levels of the integrin α4 (a subunit of the integrin VLA4, very late antigen 4), we did not find a positive correlation (**Figure S4C**). Nonetheless, we found that those patients stratified as high-risk (poor early cytological response, or minimal residual disease level ≥ 0.05% at the end of reinduction), showed a positive correlation between the mRNA levels of WDR5 and α4 subunit (**Figure S4D**). To further validate the effect of WDR5 on the cell migration mediated by integrins, we observed that WDR5 silencing or inhibition did not affect the expression of α-subunit of the integrin VLA4 at the membrane of ALL cell lines (**Figures 4C and 4D)**. To further validate the effect of WDR5 activity on the cell migration mediated by integrins, we determined the activation stage of the β-subunit of the integrin VLA4 and found only a slight reduction in WDR5-depleted cells and no differences upon OICR-treatment (**Figures S4E and S4F**). More importantly, we confirmed that OICR-treatment did not affect the VLA4-dependent migration in ALL cell lines (**Figure 4E**). These results indicated that WDR5 expression correlated with cell adhesive molecules in ALL cells, and had no clear contribution on the migration mediated by these receptors. This supports our hypothesis that targeting WDR5 might impede the dissemination of ALL cells by a different mechanism.

**Fig. 5.**
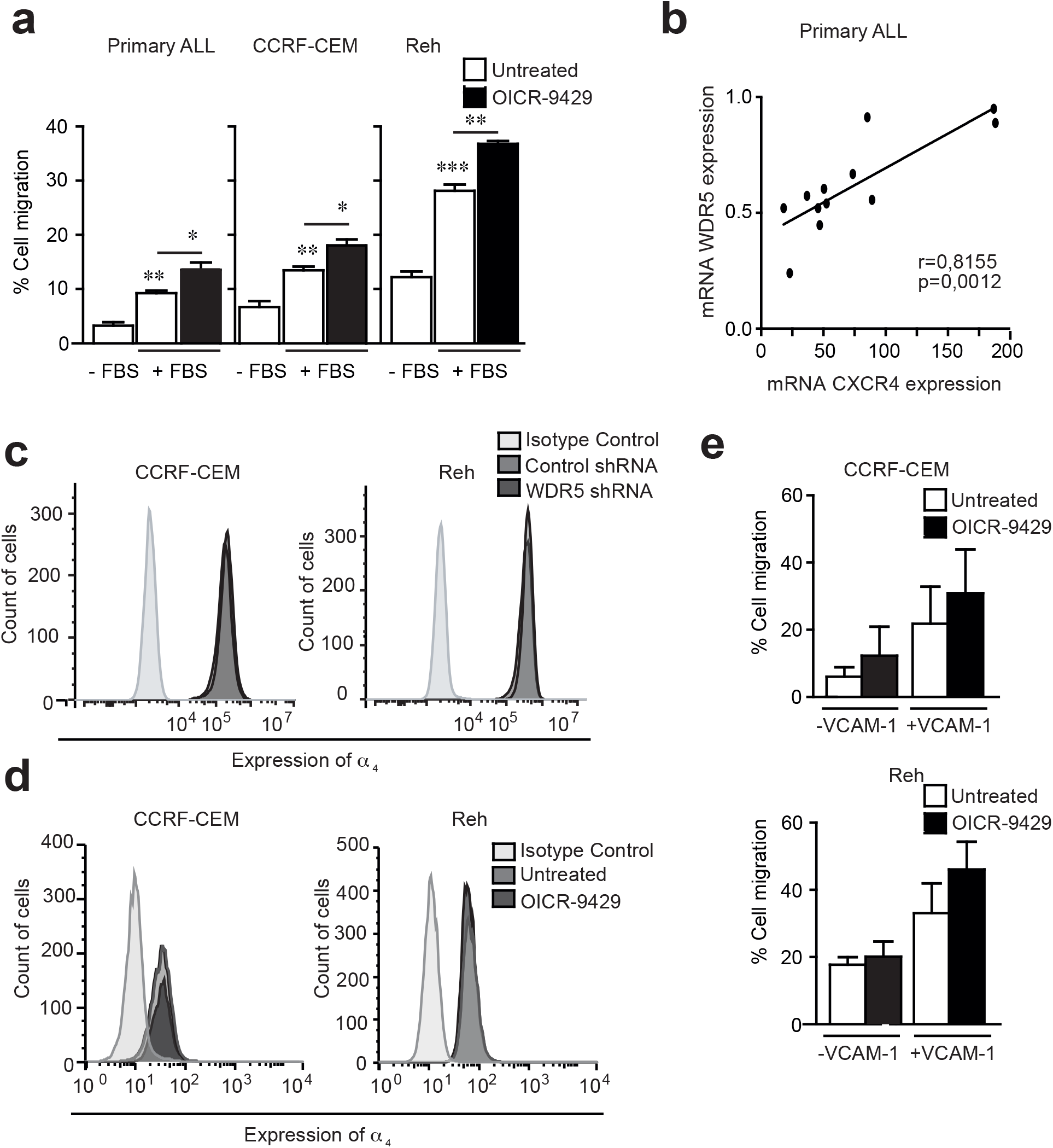
WDR5 expression associates with cell migration molecules in clinical leukemia samples. **(A)** ALL cells pretreated or not with 1 μM OICR-9429 were seeded on the top of Transwell chambers and allowed them to migrate in response to serum. Cells were collected from the bottom chamber after 24 h and quantified. Mean n = 3 ± SEM. **(B)** WDR5 and CXCR4 expression (mRNA) in ALL cells from patients (n=12) analyzed by RT-qPCR. Expression levels were normalized by TBP and the graph shows the correlation between both molecules. Pearson’s correlation coefficient (r) and P-value are shown. **(C)** Flow cytometry expression of α4 subunit at the surface of CCRF-CEM and Reh cells transfected with control or WDR5 shRNAs. **(D)** Flow cytometry expression of α4 subunit at the surface of CCRF-CEM and Reh cells pretreated or not with 1 μM OICR-9429. **(E)** CCRF-CEM and Reh cells pretreated or not with 1 µM OICR-9429 were seeded on the top of Transwell chambers coated with VCAM-1 (10 µg/ml). Cells were collected from the bottom chamber after 24 h and spontaneous migration induced by VCAM-1 adhesion quantified. Mean n = 4 ± SEM. (*) P < 0.05, (**) P < 0.01, (***) P < 0.001.

Leukemia cell migration depends on integrins and chemokine receptors,^3^ and impaired ALL infiltration might be related to defects on how ALL cells respond to chemotactic gradients. Interestingly, OICR-9429 treatment of primary samples and ALL cell lines did not reduce the migration towards chemoattractant (**Figure 4A)**. We also confirmed that WDR5 depletion did not reduce the chemotactic migration of CCRF-CEM and Reh cells (**Figure S4A**). To determine how WDR5 expression might be linked to clinical features and adhesive molecules, we analyzed the expression of WDR5 in primary samples from ALL patients. We found that WDR5 levels were not related with age or gender of patients with ALL, although we observed significant differences in those patients with more than 80% of blast in the bone marrow at diagnosis (**Figure S4B**). We found a positive correlation between the mRNA levels of WDR5 and the receptor CXCR4 (C-X-C Motif Chemokine Receptor 4) in primary samples from patients with ALL (**Figure 4B**). When we compared WDR5 expression and the mRNA levels of the integrin α4 (a subunit of the integrin VLA4, very late antigen 4), we did not find a positive correlation (**Figure S4C**). Nonetheless, we found that those patients stratified as high-risk (poor early cytological response, or minimal residual disease level ≥ 0.05% at the end of reinduction), showed a positive correlation between the mRNA levels of WDR5 and α4 subunit (**Figure S4D**). To further validate the effect of WDR5 on the cell migration mediated by integrins, we observed that WDR5 silencing or inhibition did not affect the expression of α-subunit of the integrin VLA4 at the membrane of ALL cell lines (**Figures 4C and 4D)**. To further validate the effect of WDR5 activity on the cell migration mediated by integrins, we determined the activation stage of the β-subunit of the integrin VLA4, and found only a slightly reduction in WDR5-depleted cells and no differences upon OICR-treatment (**Figures S4E and S4F**). More importantly, we confirmed that OICR-treatment did not affect the VLA4-dependent migration in ALL cell lines (**Figure 4E**). Collectively, our data suggest that WDR5 expression might correlate with cell adhesive molecules in leukemia cells.

### H3K4 methylation induced by 3D conditions is dependent of actomyosin cytoskeleton and regulates the cell motility through confined conditions

Cell motility in 3D conditions is independent of integrin signaling and requires highly nuclear deformability and mechanotransduction.^17^ Then, we first interrogated how leukemia cells move in 3D confined conditions. We quantified the spontaneous displacements of CCRF-CEM cells embedded in a 3D collagen matrix by live-cell imaging (**Movie S1**). Our quantitative analysis showed the trajectories of cell migration in 3D environments (**Figure 5A**). Importantly, we observed that the inhibition of WDR5-MLL interaction significantly reduced the speed and net displacement of CCRF-CEM cells migrating in 3D confined conditions (**Figures 5B and 5C**). We also confirmed the inhibitory effect of OICR-9429 on the migration of Reh cells embedded in a 3D collagen matrix (**Movie S2 and Figures S5A-C**). The cell cytoskeleton acts as a mechanotransducer to integrate external 3D stimuli from the cell surface into the nucleus and might be a potential therapeutic target to improve cancer therapy,^18^ therefore we addressed whether targeting WDR5 might affect the cytoskeleton of ALL cells. Flow cytometry showed that depletion or inhibition of WDR5 did not affect the level of F-actin in CCRF-CEM and Reh cells (**Figures 5D and S5D**). Furthermore, actomyosin is critical for H3K4 methylation induced by WDR5.^19^ Mechanistically, we confirmed that MLCK (myosin light chain kinase) activity was required for the H3K4 methylation induced by 3D conditions in CCRF-CEM and Reh cells, as its specific inhibitor (ML-7) impaired the upregulation of H3K4me3 levels (**Figures 5E**). We determined that MLCK localized in those chromatin regions enriched for H3K4 methylation in CCRF-CEM and Reh cells (**Figures 5F**), and found remarkable levels of this protein in the nuclear fractions of ALL cell lines and primary ALL cells (**Figures S5E**). Collectively, our data established that targeting WDR5 diminished the capacity of ALL cells to move across 3D barriers and that MLCK activity was a molecular mechanism operating downstream of this external stimulus.

**Fig. 5.**
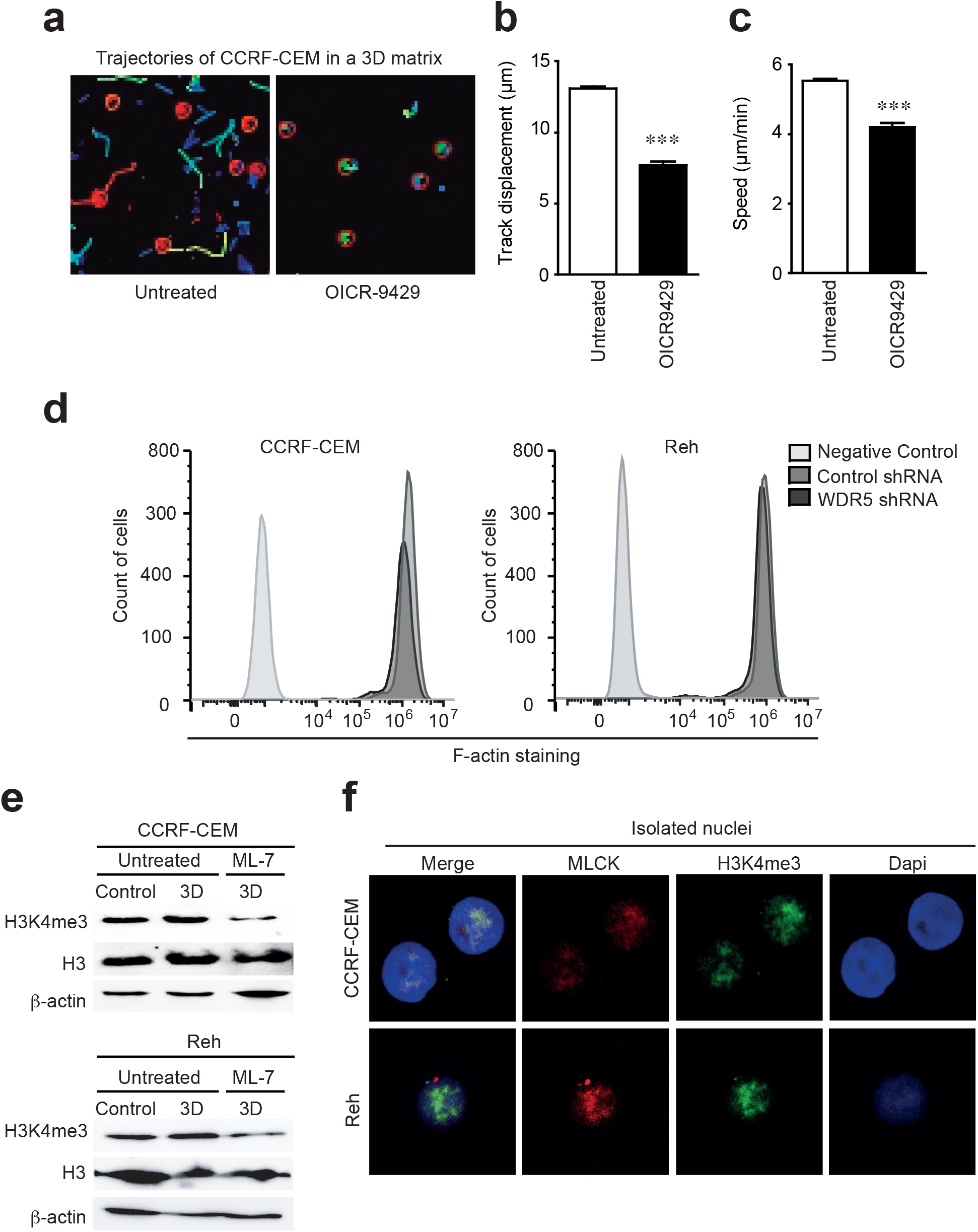
H3K4 methylation induced by 3D conditions depends on MLCK activity and controls ALL cell displacement in confined conditions. **(A)** CCRF-CEM cells were pretreated or not with 1 μM OICR-9429 (WDR5 inhibitor) for 1h, embedded in a collagen gel and allowed to migrate randomly for 3h. Cell trajectories were tracked and indicated. **(B)** Graph shows the mean of the track displacement from CCRF-CEM cells migrating in 3D conditions as in (A). Mean n = 3 independent experiments ± SEM. **(C)** Graph shows the mean of the speed of CCRF-CEM cells migrating in 3D conditions as in (A). Mean n = 3 independent experiments ± SEM. **(D)** CCRF-CEM and Reh cells transfected with control or WDR5 shRNAs were fixed and actin polymerization was determined by flow cytometry. **(E)** CCRF-CEM and Reh cells were treated with 1 µM ML-7 (MLCK inhibitor) for 30 min prior their culture in 3D conditions. After 1 h cells were collected and H3K4me3 levels tested by western blotting. **(F)** Isolated nuclei from CCRF-CEM, Reh, and primary ALL cells were seeded on polylysine-coated coverslips. Samples were fixed, permeabilized and the middle section of the nucleus was analyzed by confocal microscopy using specific antibodies. (***) P < 0.001.

### 3D culture conditions facilitate chromatin accessibility and nuclear mechanics of leukemia cells

As OICR-9429 treatment did not impair ALL cell migration upon integrin and chemokine stimulation, we focused on the role of the nucleus for cell squeezing through narrow spaces. Firstly, we sought to gain insights into how H3K4 methylation induced by 3D environmental conditions might regulate the global chromatin conformation in ALL cells. We analyzed the chromatin accessibility of CCRF-CEM and Reh cells in suspension or embedded in 3D collagen gels. An open chromatin conformation due to 3D conditions was associated with increased chromatin accessibility to micrococcal DNAse (MNAse) digestion (**Figure 6A**). We observed that 3D conditions promoted more sensitivity to DNAse digestion and bigger open peaks of nucleosome releasing (**Figure 6B**). To complement our data and confirm morphological and chromatin changes in the nucleus of ALL cells embedded in 3D conditions, we used electron microscopy to image CCRF-CEM cells cultured in suspension and in 3D conditions. We visualized more rounded and bigger nuclei corresponding to cells cultured in 3D conditions, compared to cells in suspension (**Figure 6C**). Then, we tested in isolated nuclei from primary ALL cells how 3D constricted conditions might regulate the mechanical response of the nucleus upon external forces (**Figure 6D)**. We quantified the nuclear area pre-and post-confinement and determined increased nuclear area and deformability of isolated nuclei from primary ALL cells in 3D conditions (**Figure 6E)**. To determine the fundamental impact of H3K4 methylation on the mechanical changes induced by 3D conditions, we cultured control and OICR-9429-treated cells in suspension or embedded in 3D environments. We confirmed that WDR5 inhibition reduced the effect of 3D conditions (**Figure 6F**). Similarly, WDR5 depletion also abrogated the increased nuclear deformability induced by 3D conditions (**Figure S6**). Together, these data suggested that H3K4 methylation induced by 3D constrained conditions altered the global chromatin structure and mechanical properties of ALL cells.

**Fig. 6.**
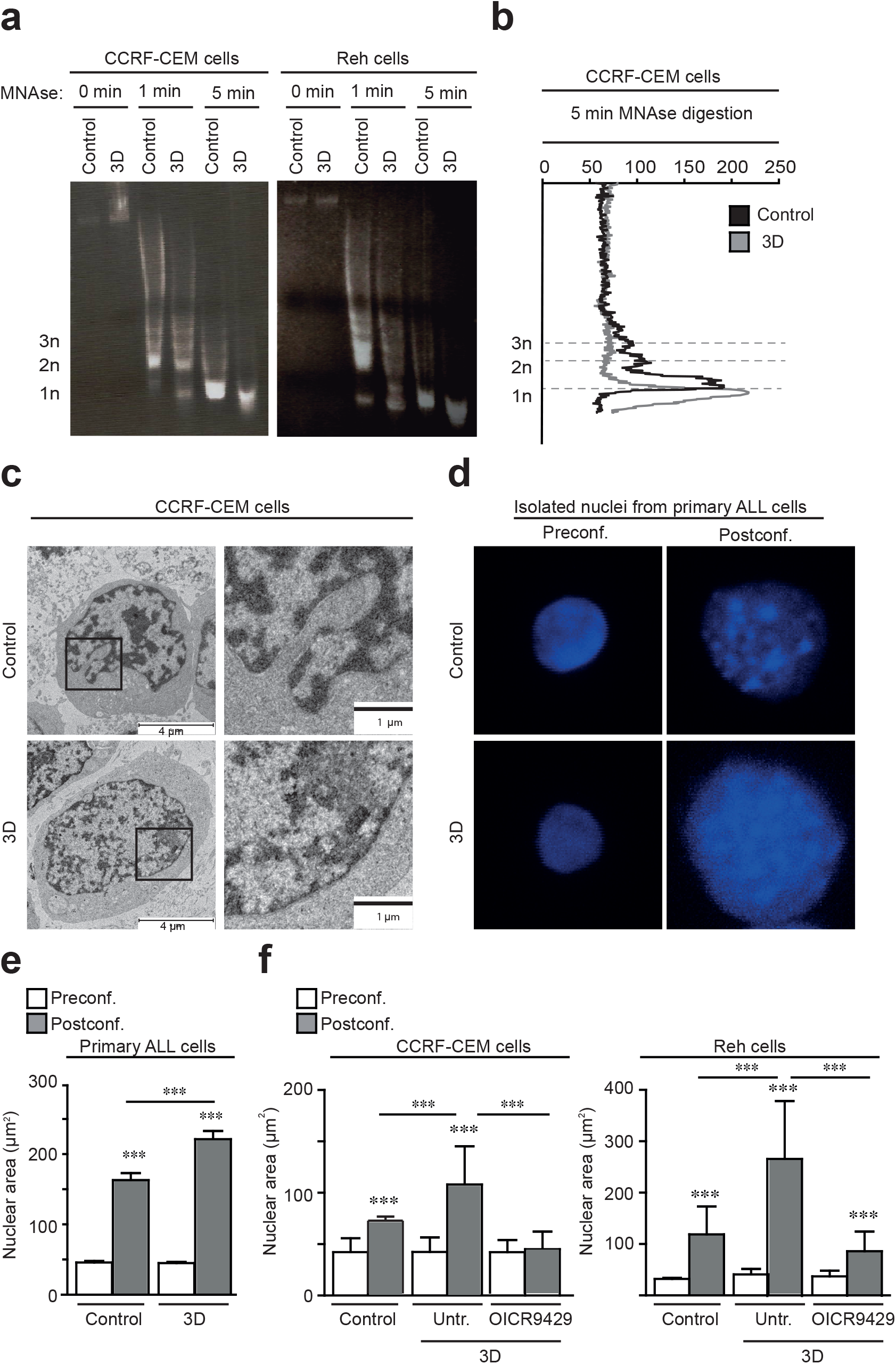
3D confined conditions promote conformational and mechanical changes in the chromatin and the nucleus of leukemia cells. **(A)** CCRF-CEM and Reh cells were cultured suspension (Control) or embedded in a 3D collagen matrix (3D). Then, cells were collected and digested with micrococcal nuclease at indicated times. DNA fragments were purified and resolved in agarose gel. Mono (1n), di (2n) and tri (3n) nucleosomes are indicated. **(B)** Graph shows the nucleosomal releasing profile from CCRF-CEM cells at 5 min after micrococcal digestion. Mono (1n), di (2n) and tri (3n) nucleosomes are indicated. **(C)** Electron microscopy of CCRF-CEM cells cultured in suspension or embedded in a 3D collagen matrix. Black squares on the left panel represent the area selected for detail on the right panel. **(D)** Isolated nuclei from primary ALL cells were stained and seeded on polylysine-coated coverslips. Confocal sections of the nuclei were taken before (Preconf.) and after (Postconf.) confinement. **(E)** Graph shows the mean average of the nuclear area in (D). Mean n = 22-46 isolated nuclei ± SEM. **(F)** CCRF-CEM and Reh cells were treated with 1 μM OICR-9294 for 1h prior to their culture in 3D conditions. Then, their nuclei were isolated, confined and their nuclear area was quantified. Mean n = 53-71 isolated nuclei ± SEM. (***) P < 0.001.

## Discussion

The cell migration through 3D environments is a critical step in multiple physiological and pathological processes such as development, inflammation, wound healing, and cancer invasion.^5^ The cell movement in these confined conditions constricts the nucleus, which limits the capacity of cells to cross physical barriers.^17^ To penetrate into protective niches, such as the bone marrow or the central nervous system, cancer cells must respond to biochemical and mechanical signals induced by their surrounding 3D environment.^20,21^ Therefore, identifying novel molecules that regulate molecular and mechanical pathways for tumor dissemination is an important direction to developing more effective therapeutic options. In this study, we have defined a mechanism used by ALL cells to promote H3K4 methylation and alter the mechanical properties of their nuclei during cell infiltration in 3D environments.

3D culture conditions lead to transcriptional and epigenetic changes, altering the ratio between hetero and euchromatin.^22,23^ Firstly, we determined that 3D confined conditions induced upregulation of H3K4me3 of ALL cells. Mechanistically, the actin dynamics and myosin contractility translate intracellular signals and forces critical for hematopoiesis,^24^ leukemia infiltration,^25^ and for nuclear changes during the cell migration through complex environments;^26^ and we have demonstrated that 3D conditions promoted H3K4 methylation in ALL cells through MLCK activity. Confirming the functional link between actomyosin contractility and WDR5/H3K4 methylation,^19^ we observed that MLCK localized in the vicinity of H3K4me3 regions in the nucleus of ALL cells. This also agrees with previous observations of MLCK and myosin in the nucleus of several cell types,^27^ which suggests a potential role of this kinase in chromatin homeostasis and conformation.

It is well known that H3K4me3 is an activating histone modification linked to gene transcription,^28^ which in turn regulates the fate and behavior of the cells.^29^ Furthermore, WDR5 interacts with the protooncogene myc and the transcription factor C/EBPa to drive tumorigenesis and cancer progression;^30,31^ nonetheless, we did not find significant changes in the expression of c-Myc at the studied times. We found that 3D culture-mediated H3K4 methylation promoted genome-wide and transcriptional changes in ALL cells, suggesting that the whole nucleus might be affected during the infiltration process. Our genomic analysis showed epigenetic and transcriptional regulation of genes related to DNA repair and cell cycle. According to our results, it has been reported that WDR5 regulates the expression of genes involved in DNA repair and damage in leukemia cells.^32^ These transcriptional changes might be related to microenvironmental signals that promote clonal genetic differences, chromatin instability and cell resistance to drug therapy at specific niches.^33^ Interestingly, there was no correlation between our H3K4me3 ChIPseq and the transcriptional analyses. This discrepancy has been reported before for other histone methylations,^34,35^ and it might indicate that H3K4 methylation might have additional functions rather than transcriptional activation.

Epigenetic changes are fundamental to defining new biomarkers and potential therapeutic targets in ALL.^7,36^ We found that mRNA levels of WDR5 correlated with CXCR4 and the integrin VLA4, especially in those patients stratified as high-risk. Those cell receptors are critical for ALL cell invasiveness, and their expression has been linked to worse prognosis and disease progression;^37,38^ and VLA4 expression is linked to other histone methyltransferases.^39^ This suggests that there might be a functional link between WDR5 and surface cell receptors, and that the expression of these molecules might serve to promote ALL cell invasion at different stages.

Histone methylation has been proposed as a potential therapeutic target in cancer.^40^ It has been reported that OICR-9429 treatment blocks cancer cell proliferation and growth.^41^ We did not observe significant differences in the cell cycle and DNA damage in ALL cells upon OICR-9429 treatment, which might be due to the short times analyzed to distinguish how H3K4 methylation might control cell migration without affecting other cellular functions. By using a murine in vivo model to study ALL spreading through the whole organism, we confirmed that WDR5 activity improved ALL cell invasiveness and a migratory benefit to colonizing the spleen and the bone marrow. ALL infiltration into collagen matrices might be regulated by chemokine receptors, which have been reported critical for the migration and invasiveness of ALL cells.^43^ Although targeting WDR5 severely reduced cell migration across 3D environments and *in vivo* invasiveness, we did not observe that blocking WDR5 activity reduces the chemotaxis of ALL cells. Furthermore, actin polymerization (F-actin) is known to control adhesive structures related to cancer cell migration and adhesion,^44^ but our data suggested that the inhibitory effect of WDR5 inhibition on ALL cells was not mediated by affecting actin polymerization, nor the expression of the integrin VLA4 in ALL cells. In accordance with this finding, it has been reported that WDR5 regulates the invasive properties of breast cancer cells without inhibiting cell adhesion.^14^

To determine the biomechanical mechanism used by leukemia cells to migrate across confined environments, we focused on the role of the nucleus. Nuclear deformability has emerged as a master regulator of cancer cell migration.^44^ It is well described that the mechanical properties of the nucleus are defined by the nuclear lamina and the chromatin structure.^45,46^ Several histone methylations are promoted during cell migration and physical stress.^34,47^ We confirmed that isolated nuclei from ALL cells cultured in 3D showed higher nuclear deformability than cells in suspension, and OICR-9429 treatment impaired the mechanical change induced by 3D conditions. Interestingly, it has been reported that the loss of H3K4me3 has minor effects on transcription and other functional effects such as DNA replication stress,^48-50^ which suggest that H3K4 methylation might have other non-genomic functions that might confer a significant advance for ALL cells to infiltrate other tissue and organs.

In summary, our study indicated a novel nuclear mechanism used by ALL cells to infiltrate constrained 3D tissue spaces. This mechanism evolves rapid dynamic adaptation of H3K4 methylation to control the nuclear mechanical properties and transcriptional changes in ALL cells. Our findings highlight a relevant function of H3K4 methylation for ALL cell infiltration and plasticity that might provide therapeutic possibilities against leukemia dissemination.

## Acknowledgements

The authors thank the Microscopy Unit of Instituto de Investigación Biosanitaria Gregorio Marañón (IiSGM) for assistance with confocal and videomicroscopy analyses. The authors are also grateful to the EM and Animal Facilities of platforms of the CIB Margarita Salas for their assistance and support with the EM and *in vivo* experiments. The UCM-Genomic CAI Unit for their assistance with microarray experiments. This research was supported by a FPI Scholarship 2018 (Ministerio de Ciencia e Innovacion/MICINN, Agencia Estatal de Investigacion/AEI y Fondo Europeo de Desarrollo Regional/FEDER) to R.G.N.; grants from Asociación Pablo Ugarte to M.R.; and grants from 2020 Leonardo Grant for Researchers and Cultural Creators (BBVA Foundation), Ayuda de contratación de ayudante de investigación PEJ-2020-AI/BMD-19152 (Comunidad de Madrid), and the Ministerio de Ciencia e Innovacion (MICINN) Agencia Estatal de Investigacion (AEI) y Fondo Europeo de Desarrollo Regional (FEDER) (SAF2017-86327-R) to J.R.M.

## Authors’ contributions

R.G.N., A.dL.P., A.G.M., D.A., E.M., conducted experiments. M.R., provided primary ALL samples, clinical characterization and contributed to clinical data interpretation. M.R., J.R.M designed and supervised the experiments, wrote the paper, and provided financing.

## Ethics approval and consent to participate

ALL primary cells were obtained from patients under 16 years old with informed consent for research purposes and approved by the Ethics Committee for Medical Research (IRB) at Hospital Universitario Niño Jesús and CSIC. All experiments involving animals were approved by the OEBA (Organ for Evaluating Animal Wellbeing) at CIB Margarita Salas and Madrid Regional Department of Environment, with reference PROEX 228.4/21.

## Data Availability Statement

Genome-wide sequencing raw reads and processed files have been deposited at Gene Expression Omnibus (GEO). The accession numbers for the ChIP-seq and the transcriptional microarray data are GSE193383 and GSE190771, respectively. All datasets, materials, and software used in this study are listed in Methods and Additional files.

