## Supplemental Info for "Targeting H3K4 methylation as a novel therapeutic strategy against tumor infiltration and nuclear changes of acute lymphoblastic leukemia cells"

**Supplementary Materials and Methods**

**Supplementary Figures**

**Supplementary Tables**

**Supplementary Movies**

| **Antibodies and Reagents** | **Host** | **Company** | **# Cat** |
|---|---|---|---|
| anti-H3K4me3 | Rabbit | Cell signaling | 9751S |
| anti-H3K27me3 | Rabbit | Cell signaling | 9733 |
| anti-H3 total | Rabbit | Cell signaling | 4499 |
| anti-H3K9me2/3 | Mouse | Cell signaling | 5327 |
| anti-H3K9ac | Rabbit | Upstate | 06-599 |
| anti-Cyclin D1 | Mouse | Sigma-Aldrich | C7464 |
| anti-β-actin | Mouse | Sigma-Aldrich | A1978 |
| anti-MLCK | Rabbit | Sigma-Aldrich | SAB1300116 |
| anti-Lamin B1 (C-5) | Mouse | Abcam | sc-365962 |
| anti-c-myc | Rabbit | Santacruz | sc-40 |
| anti-WDR5 (G-9) | Mouse | Santacruz | sc-393080 |
| anti-RhoGDI-α (A-20) | Rabbit | Santacruz | sc-360 |
| anti-rabbit IgG (H+L). Dylight 680 | Donkey | Invitrogen | SA5-10042 |
| anti-mouse IgG (H+L). Dylight 800 | Donkey | Invitrogen | SA5-10172 |
| anti- pH2AX | Mouse | Invitrogen | 14-9865-82 |
| anti-rabbit HRP- conjugated | Goat | BioRad | 170-6515 |
| anti-mouse HRP- conjugated | Goat | BioRad | 170-6516 |
| CF647- anti-mouse IgG (H+L) | Donkey | Biotium | 20042 |
| CF594- anti-rabbit IgG (H+L) | Donkey | Biotium | 20152 |
| CF488A- anti-mouse IgG (H+L) | Donkey | Biotium | 20014 |
| Phalloidin-CF488A | | Biotium | 42 |
| α4 (HP2/1) | Mouse | Prof Sánchez-Madrid | - |
| ß1 (HUTS-21) | Mouse | Prof Sánchez-Madrid | - |
| Dapi | | Thermo Scientific | 62248 |
| CellTrace™ CFSE | | Thermo Scientific | C34554 |
| CellTrace™ Far Red | | Thermo Scientific | C34564A |
| BCECF-AM | | Thermo Scientific | C34564A |
| Bovine collagen solution type I | | Stem Cell | 7001 |
| Human VCAM-1 | | Peprotech | 150-04 |
| OICR-9429 | | Tocris Bioscience | 5267 |
| Hoechst 33258 | | Sigma-Aldrich | 94403 |
| Propidium iodide | | Sigma-Aldrich | P4864 |
| ML-7 | | Sigma-Aldrich | 110448-33-4 |
| Poly-L-Lysine | | Sigma-Aldrich | 25988-63-0 |
| DACO | | Sigma-Aldrich | 10981 |
| Proteinase K | | Sigma-Aldrich | P1308 |
| RNAse | | Sigma-Aldrich | R4875 |
| rProtein A sepharose | | Sigma-Aldrich | P9424 |
| 35mm glass-bottomed plates | | Ibidi | 80136 |
| Micrococcal nuclease | | New England Biolabs | M0247S |
| Phenol:chloroform:isoamyl alcohol | | Panreac | A0889.0250 |

**H3K4me3 quantification and Immunoblotting**. Cells were washed with cold PBS, lysed in sample buffer (VWR) and then sonicated (15 s at 70% amp) using a Microson XL2000 (Misonix) to extract the histones. For cellular fractionation, we followed the reported Wung P 2017 (PNAS) protocol. The cytoplasmic fraction was extracted in buffer A (10 mM HEPES, 10 mM KCl, 1.5 mM $MgCl_2$, 0.34 M sucrose, 10% (v/v) glycerol, 1 mM DTT and Roche protease inhibitor) for 5 min at 4ºC and centrifuged for 5 min at 4°C at 3,500 g. Then, cytoplasmic fractions were collected from centrifugation at 3200 rpm for 5 min. Pellets were resuspended in RIPA buffer (NaCl 1M, NP-40 1%, Sodium deoxycholate 0.5%, SDS 0.1%, Tris-HCl [pH 7.4] 50mM) containing protease inhibitors for 5 min. After 15 min, soluble nuclear fractions were collected from centrifugation (5 min, 4°C, 13500 rpm) and chromatin bound fractions were resuspended in sample buffer and sonicated. Samples were boiled and proteins separated in 7.5-15% polyacrylamide gels. They were transferred to nitrocellulose membranes (GE Healthcare Life Science) that were sequentially blocked in 5% low fat milk in TBS-Tween (0.5%) for 1 hour at room temperature and incubated overnight with primary antibodies at 4ºC with rotation. After washing with TBS-Tween 1%, membranes were incubated with secondary antibodies in 5% low fat milk in TBS-Tween (0.5%) for 1 hour at room temperature and protein signal was resolved.

**Transwell invasion.** $2x10^5$ cells under specific conditions were added on the upper chamber of transwell inserts (Corning Costar, 6.5 mm diameter, 5 µm pore sizes), and 600 µL of medium with or without serum to the bottom chamber. In some cases, cells were added onto 10 µg/mL VCAM-1 coated chambers, to analyze the migration induced by integrin attachment. After 24 h, migrated cells were collected from the bottom chamber and counted to calculate migration index.

**Flow cytometry.** ALL cells were blocked with human IgG (50 µg/ml; 30 min), incubated with primary antibodies (5-10 µg/ml; 30 min), washed twice with PBS, followed by appropriated fluorochrome-conjugated secondary antibodies for 30 min. For F-actin quantification, cells were fixed with PFA 4% for 5 min at RT, permeabilized with PBS Triton x-100 0.1% for 5 min and stained with Phalloidin 50 µg/mL for 1h. Flow cytometry was performed with a FacSort (Beckman Coulter) and data were analyzed using the Flow Jo software (Flow Jo LLC, Ashland, OR, USA).

**MNAse digestion.** To analyze the nucleosomal profile, suspension or 3D matrix embedded cells were lysed in lysis buffer (10 mM Tris [pH 7.5], 10 mM NaCl, 2 mM MgCl2, 0.5% NP-40, 1 mM CaCl2) with protease inhibitors. 100U of MNAse were added for each time point and samples were warmed at 37ºC to activate the enzyme. Reactions were stopped by the addition of EDTA after 1 or 5 min of digestion. Samples were incubated with RNase A for 10 min at 37°C and then with 0.01% SDS and proteinase K overnight at 65°C. Extraction with 1 volume of phenol-chloroform-isoamyl alcohol and precipitation with 0.1 volumes of sodium acetate (3M) with 2.5 volumes of ethanol overnight at -20ºC were performed. Samples were high speed centrifuged for 30 min at 4ºC and the DNA pellet was dissolved in water. Nucleosomal releasing was resolved in 2% agarose gel and DNA digestion profiles were quantified with ImageJ.

**3D migration assay.** Control or pretreated cells were labeled with 1 µM Cell Tracer Far Red or 5 µM CFSE vital-dyes. Then, cells were washed and both populations mixed at 1:1 prior embedded in a 3D collagen matrix. Random cell migration was analyzed from images taken every 5 min during 2 h. Cell migration was tracked and analyzed with National Institutes of Health (NIH) ImageJ TrackMate plug-in.

**Nuclear confinement**. ALL cells were cultured in suspension or embedded in a 3D matrix for 1 hour and their nuclei were isolated using buffer A (10 mM HEPES, 10 mM KCl, 1.5 mM

$MgCl_2$, 0.34 M sucrose, 10% (v/v) glycerol, 1 mM DTT and Roche protease inhibitor). Nuclei were resuspended in TKMC buffer (50 mM Tris pH 7.5, 25 mM KCl, 3 mM MgCl2, 3 mM CaCl2, and proteinase inhibitors) and dyed with DAPI 1 μg/mL. Then, nuclei were sedimented onto poly-L-lysine coated plates and placed in the cell confiner (4D cell) following the manufacturer's instructions. In this case, a glass slide with micropillars of 3 μm height was stuck on the silicone macropillar attached to the lid of the confiner. When the lid was closed, the pillars pushed the confining slides onto the culture substrate and confined the nuclei to 3 μm. Images of at least 20 nuclei were taken with a 63× objective before and after the confinement by an inverted confocal microscope DMi8 (Leica) and their 3D reconstructions were performed with Leica software (LCS-1537). The nuclear area was quantified using ImageJ software.

**Chromatin Immunoprecipitation-Sequencing (ChIP-Seq).** CCRF-CEM cells were cultured in suspension or embedded in a 3D collagen matrix with RPMI for 1h at 37ºC 5% $CO_2$. After washing and straining, cells were fixed with paraformaldehyde 1% for 4 min at room temperature. The crosslinking was stopped with 0.125M glycine during 5 min at RT. Cells were washed with PBS, centrifuged at 1800 rpm 2 min 4ºC and sonicated in cell membrane lysis buffer (1% SDS, 1 mM EDTA, 10 mM Tris–HCl, pH 8.1) using a Diagenode Bioruptor (10 cycles 30s ON/30s OFF) and maintaining temperature below 20ºC. The chromatin (supernatant) was obtained by high-speed centrifugation for 10 min at 4ºC. From 25 μL of chromatin, DNA was isolated and loaded in a 1.5% agarose gel to check that the DNA fragments' size was between 100-400 bp. The rest was frozen at -80ºC overnight. 3 μg of chromatin resuspended in RIPA buffer were incubated with 60 μL of rProtein A sepharose with rotation for 30 min at 4ºC. Samples were centrifuged at 1000 rpm for 1 min and 3 μL of rabbit anti-H3K4me3 antibody was added to the supernatant and incubated for 3h at 4ºC on rotation. Immunoprecipitation was performed by addition of 60 μL of rProtein A sepharose and incubation overnight with orbital movement at 4ºC followed by 3 washes with Low salt wash buffer (0.1% SDS, 1% Triton x-100, 2mM EDTA, 20 mM Tris pH 8.0, 150 mM NaCl), Medium Salt Wash buffer (1% NP-40, 1mM EDTA, 10mM Tris HCl pH 8, 250 mM NaCl) and High Salt Wash Buffer (0.1% SDS, 1% triton x-100, 2mM EDTA, 20 mM Tris pH 8.0, 500 mM NaCl). Protein A-chromatin complexes were eluted with 1% SDS, 100 mM NaHCO3 and warmed to 30ºC for 15 min in a shaker. After 2000 rpm for 1 min centrifugation, the supernatant (DNA-protein complexes) was collected and reverse-crosslinked, and digestion with 2 µL of Proteinase K at 65 ºC for 2h and 1 µL of RNAse A at 37ºC for 15 min was performed. DNA was isolated using 1 volume of phenol:chloroform:isoamyl alcohol (25:24:1) and precipitated with 2.5 volumes of 100% ethanol and 3M sodium acetate overnight at -20ºC. Library preparation and sequencing of obtained DNA was performed by BGI Genomics. Analysis was performed at CIB-CSIC Bioinformatics and Biostatistics Unit and ChIPseq files used in this study will be deposited at the NCBI Gene Expression Omnibus (GEO).

**RNA microarray**. CCRF-CEM cells were cultured in suspension or embedded in a 3D collagen matrix with RPMI for 1h at 37ºC. Then, RNA was extracted using the NucleoSpin RNA kit (Macherey-Nagel) according to the manufacturer's instructions. RNA's purity and concentration were determined by Nanodrop measurement. 1 mg of RNA was used for microarray analysis by Human Gene Clariom S Assay (Thermo Fisher Scientific). Data were processed, normalized and log2 transformed by the UCM-Genomic CAI Unit. Analysis was performed using Transcriptome Analysis Console and Database for Annotation, Visualization and Integrated Discovery (DAVID) v6.8 and microarray data set used in this study will be deposited at GEO.

**Real-Time PCR (qPCR).** Total RNA was extracted (RNeasy kit; Qiagen, Madrid, Spain), 1.0 µg of total RNA was retrotranscribed using the SuperScript II First-Strand Synthesis

System and oligo(dT) (Invitrogen), and cDNAs were quantified using the Universal Human Probe Roche library (Roche-Diagnosis, Madrid, Spain). Primers for WDR5 (Fw: cagaaactacaaggccacacag; Rv: cttccacagtttaattgttttgtca), ITGA4 (Fw: gcgtggtacaacttgactgc; Rv: tcctcttccgctctgctg) and CXCR4 (Fw: tgtttccgtgaagaaaatgct; Rv: tgccagttaagaagatgatgga) were from Roche. Quantitative real-time PCR was performed on a Roche LightCycler® 480 in triplicate and results were normalized to TATA-binding protein expression (Roche Real Time Ready Single Assay ID 101145). Melt curve analysis was performed at the end of PCR to confirm the presence of a single, specific product. The results were expressed using the ΔΔCt method for quantification.

**shRNA transfection.** WDR5 (sc-61798-SH) or control (sc-37007) shRNA were purchased from Santa Cruz Biotechnology. Transfection of CCRF-CEM and Reh cells was facilitated by MISSION siRNA transfection reagent (Sigma-Aldrich) following the manufacturer's instructions. 24 hours after the transfection, the transfection efficiency was confirmed by western blotting.

**Cell cycle and survival.** Cell cycle was measured by the DNA content of ALL cells using PI staining. CCRF-CEM and Reh cells were cultured with or without OICR-9429 for 24 h. Then, cells were collected and fixed in ice-cold 70% (v/v) ethanol for 24 h at 4 °C. The cell pellet was collected by centrifugation at 1800 rpm, resuspended in PBS, and stained with a mixture of RNase (10 µg/mL) and PI (50 µg/mL) in sodium citrate containing 0.1% Triton X-100 for 30 min in the dark. Then, the cell pellet was collected by centrifugation at 1800 rpm, resuspended in PBS containing EDTA 2mM, and acquired in the cytometer (Cytoflex, Beckman Coulter). Data analysis was performed using the software FlowJo and the cell cycle distribution determined by G1, S, and G2/M cell populations. Cell survival upon OICR-9429 was detected by flow cytometry. CCRF-CEM and Reh cells were cultured with or without OICR-9429 for 24h. Then, cells were collected, washed in PBS and stained using the Annexin

V-FITC Apoptosis Detection kit (Immunostep) according to the manufacturer's instructions. Data analysis was performed using the software FlowJo. Negative cells for both Annexin V-FITC and PI were considered as living cells.

**Immunofluorescence**. ALL cells were cultured in suspension or in 3D conditions for 1 hour at 37ºC, then their nuclei were and sedimented onto glass slides coated with poly-L-lysine and fixed with 4% formaldehyde (10 min). After 5 min permeabilization with 0.5% Tx-100 in PBS, samples were blocked in 10% fetal bovine serum with 0.1% Tx-100 in PBS and incubated with appropriate primary antibodies for 1 h at RT, followed by several PBS washes and 1h at RT incubation with secondary antibodies. Samples were washed and mounted. Images were acquired on an inverted DMi8 microscope (Leica) using an ACS-APO 63x NA 1.30 glycerol immersion objective. Quantification and analysis of images were determined using ImageJ.

**Electron microscopy**. Cells were fixed for 1h in 3% glutaraldehyde in PBS and then washed twice with PBS. Samples were post-fixed in 1% osmium tetroxide and 0.8% potassium ferricyanide for 1h at 4ºC and washed with PBS prior to dehydration with an increasing gradient of ethanol (30%, 50%, 70%, 80%, 90% and 100%) of 10 min per step. Samples were embedded in LX112 resin and were polymerised for 48h at 60ºC. 60-80 nm sections were placed in copper grids of 75 mesh and stained with 5% uranyl acetate for 30 min and lead citrate for 4 min. Samples were viewed in a JEOL 1230 TEM and images were taken with a CMOS TVIPS 16 mp camera.

**Statistics**

Statistical analysis and comparisons were made with GraphPad Prism6. The numerical data are presented as mean ± SD or SEM. Differences between means were tested by Student's t test for two groups comparison. Where 3 or more groups were analyzed, one-way ANOVA was performed. P-values are indicated by asterisks ((*) $P < 0.05$; (**) $P < 0.01$; (***) $P < 0.001$).

**Supplementary Tables and Movies**

**Supplementary Table S1.** Clinical characteristics of ALL patients.

**Supplementary Table S2.** Transcriptional changes of CCRF-CEM cells cultured in suspension or in 3D conditions by microarray analysis. |Fold Change| > 1.4 and P-value < 0.05. n=3.

**Supplementary Movies 1 and 2. WDR5 inhibition impairs ALL cell movement through 3D collagen matrices.** Representative movies of CCRF-CEM (Movie S1) or Reh (Movie S2) migrating in 3D conditions. Untreated (labeled with Red Cell-Tracer) or OICR-9429 treated (labeled with CFSE) cells were embedded in a 3D collagen gel and tracked their migration through the collagen matrix for 3 h.

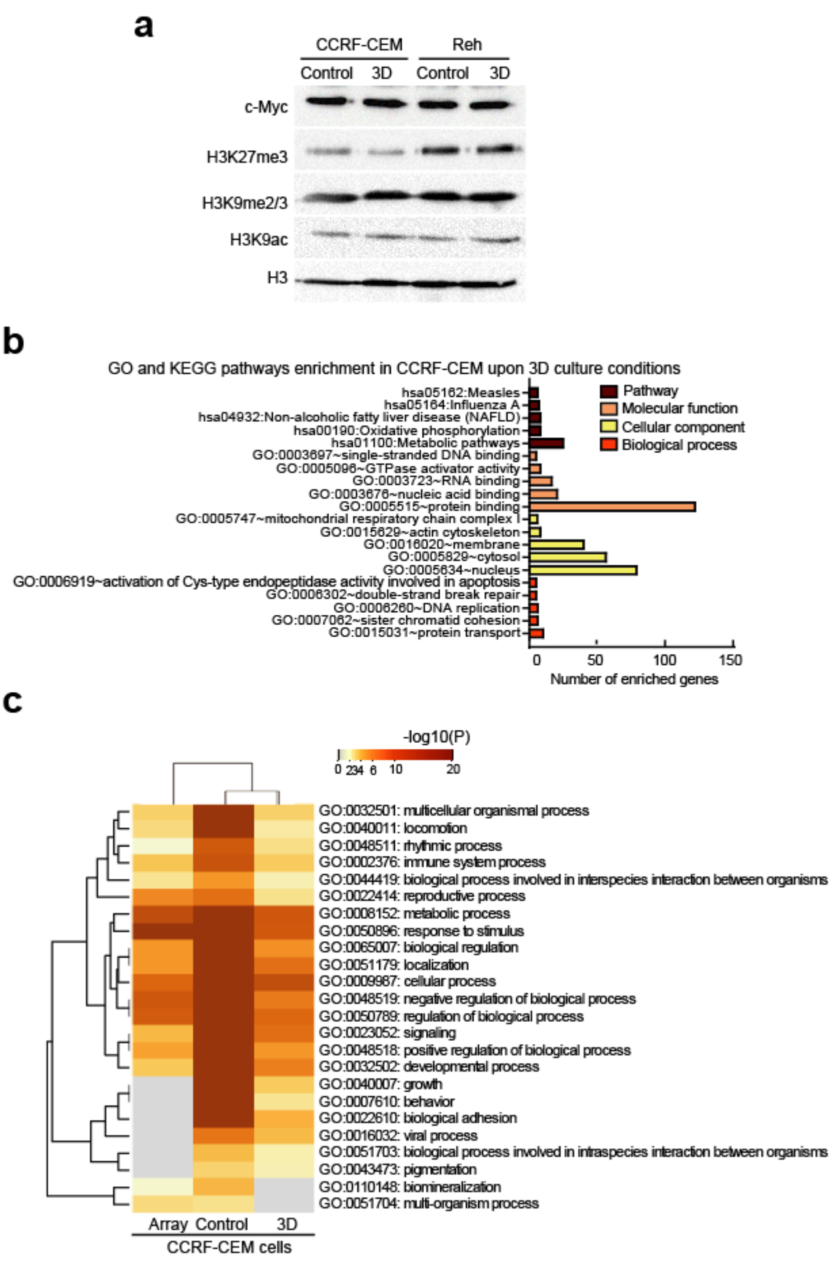


**Supplementary Fig. S1**

**Figure S1. (a)** CCRF-CEM and Reh cells were cultured in suspension (Control) or embedded in a 3D collagen matrix (3D) for 1 h. Then, the levels of different histone methylations and c-myc were tested by western blotting.**(b)** Graph shows the top 20 GO and KEGG pathways enrichment of H3K4me3 ChIPseq assay from CCRF-CEM cells cultured in suspension (control) or in a 3D collagen matrix (3D). **(c)** GO enrichment results of gene changes analyzed by mRNA microarray expression and H3K4 me3 ChIPseq analyses of CCRF-CEM cells cultured in suspension (control) and in 3D.

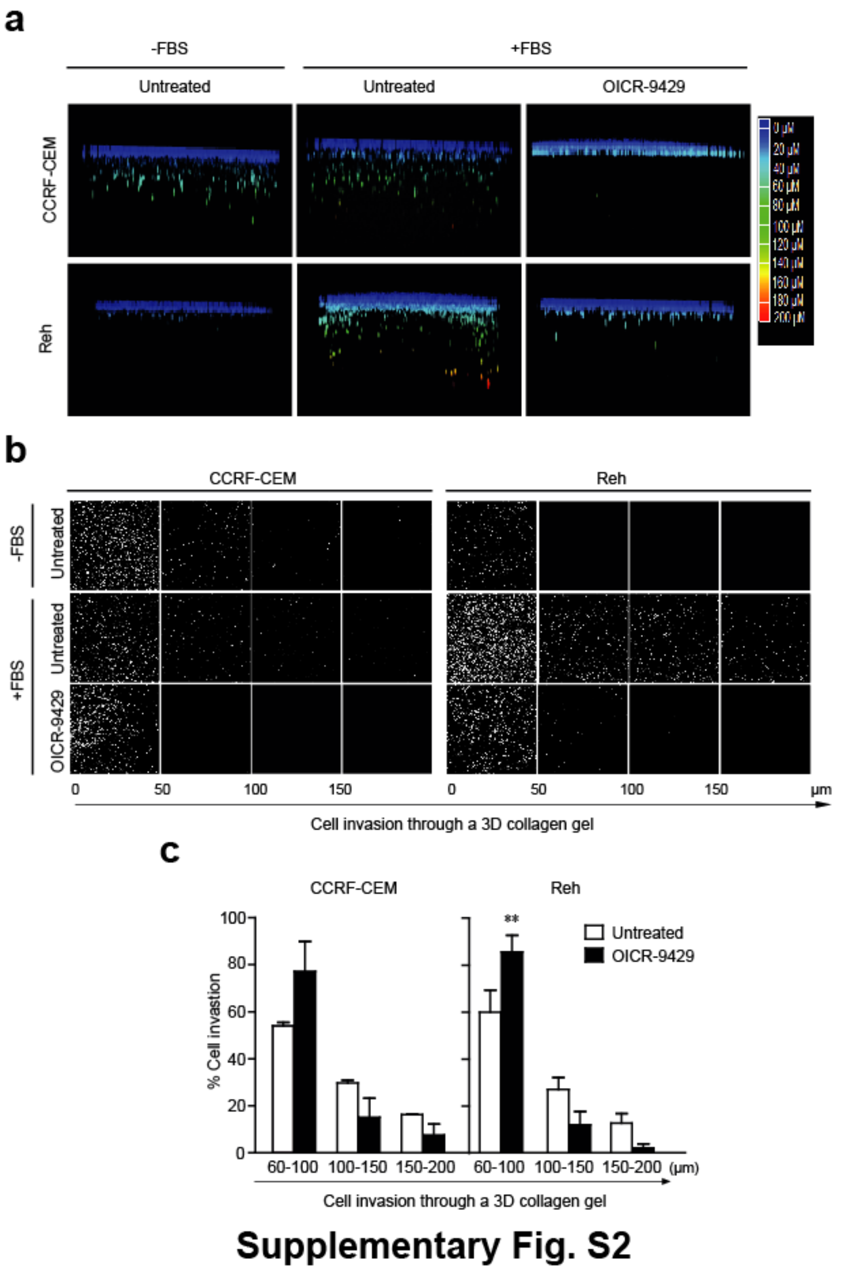


**Figure S2. (a)** CCRF-CEM and Reh cells were pretreated or not with 1 mM OICR-9429 (WDR5 inhibitor) for 1h; then, cells were seeded on the top of collagen matrix and allowed to penetrate into the collagen for 24h. Images show the cell penetrability into the 3D collagen matrix. **(b)** CCRF-CEM and Reh cells as in (A) were fixed, stained with Propidium Iodide and serial confocal sections were captured. **(c)** Graph shows the percentage of CCRF-CEM and Reh cell invasion into the collagen gel. Mean n = 3 ± SEM. (**) $P < 0.01$.

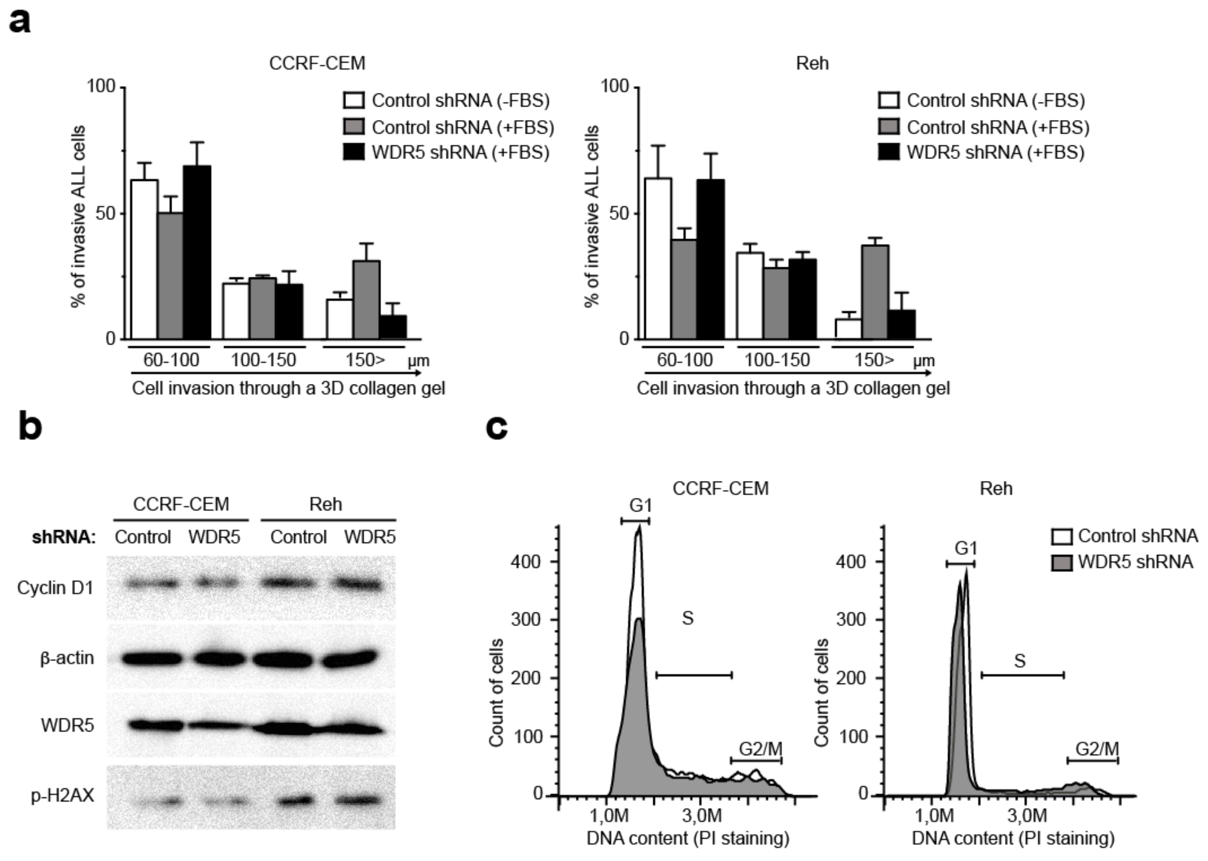


### Supplementary Fig. S3

**Figure S3. (a)** CCRF-CEM and Reh cells were transfected with control or WDR5 shRNAs for 24 h. Then, cells were seeded on the top of collagen matrix and allowed to penetrate into the collagen in response to serum (FBS, fetal bovine serum) for 24h. Cells were fixed, stained with Propidium Iodide and the percentage of cell invasion into the collagen gel was quantified. Mean n = 3 ± SEM. **(b)** CCRF-CEM and Reh cells transfected with control- and WDR5-shRNAs were lysed and protein levels analyzed by western blotting. **(c)** CCRF-CEM and Reh cells were transfected with control- and WDR5-shRNAs and their cell cycle was analyzed by flow cytometry. Graphs show the G1, S and G2/M phases according to DNA content.

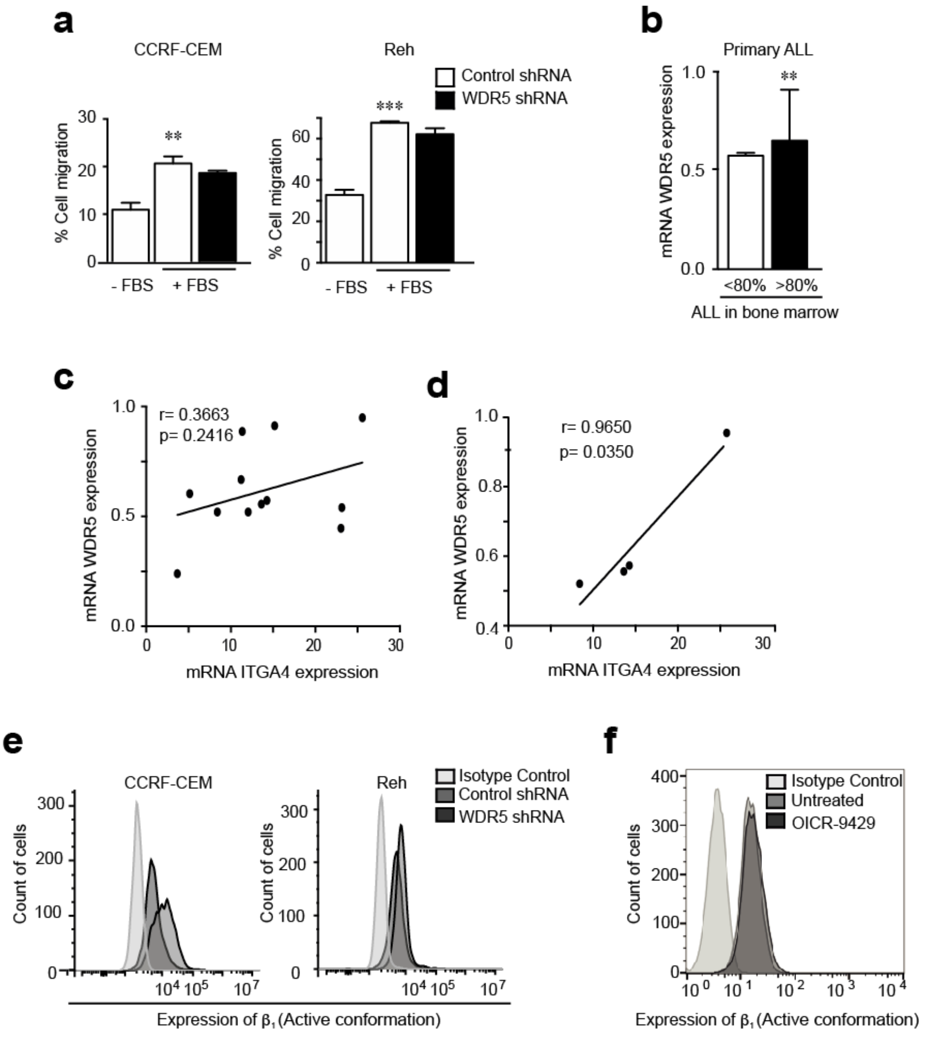


**Supplementary Fig. S4**

**Figure S4. (a)** CCRF-CEM and Reh cells were transfected with control or WDR5 shRNAs for 24 h. Graphs show the percentages of migrated cells in response to serum. **(b)** WDR5 expression (mRNA) in patients with ALL according to percentage of ALL blasts in bone marrow was determined by RT-qPCR (expression levels were normalized by TBP). Mean n=12 ± SEM. **(c)** WDR5 and ITGA4 expression (mRNA) in ALL cells from patients (n=12) analyzed by RT-qPCR. Expression levels were normalized by TBP and the graph shows the correlation between both molecules. Pearson's correlation coefficient (r) and P-value are shown. **(d)** WDR5 and ITGA4 expression (mRNA) in those patients stratified as high-risk (poor early cytological response, or MRD level ≥ 0.05% at the end of reinduction). Pearson's correlation coefficient (r) and P-value are shown. **(f)** Flow cytometry analysis of the active conformation of the VLA4 subunit β1 at the surface of CCRF-CEM and Reh cells depleted for WDR5. **(e)** CCRF-CEM cells were treated with OICR-9429 and the active conformation of the VLA4 subunit β1 integrin was determined by flow cytometry.

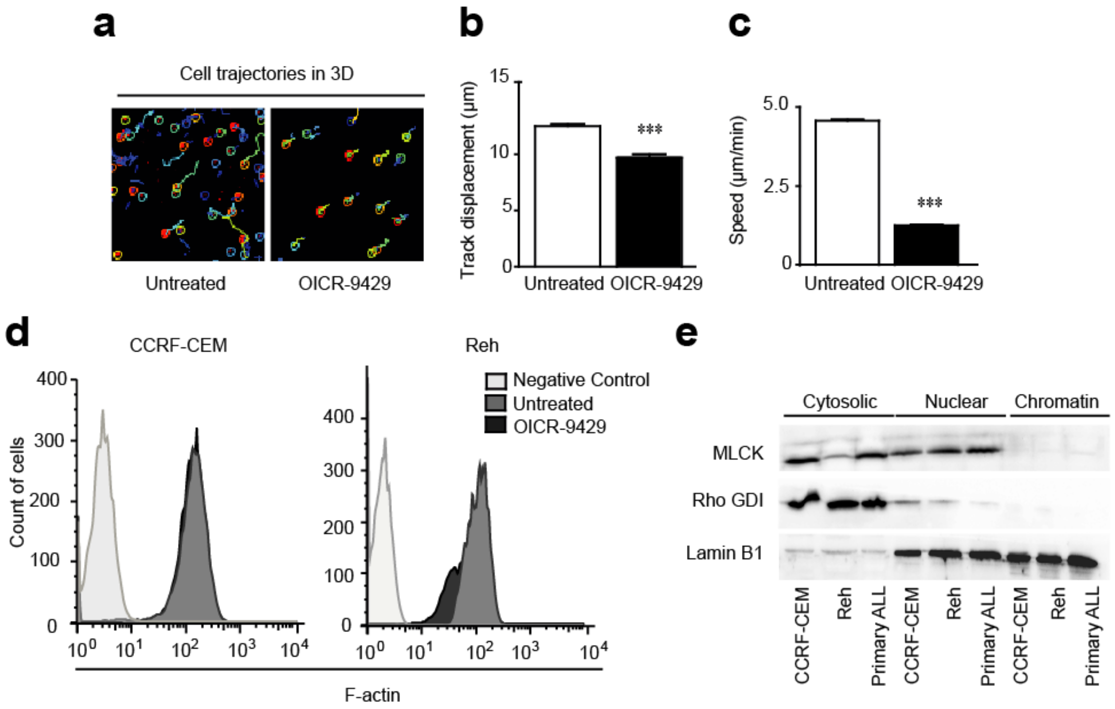


### Supplementary Fig. S5

**Figure S5. (a)** Reh cells were pretreated or not with 1 mM OICR-9429 (WDR5 inhibitor) for 1h, embedded in a collagen gel and allowed to migrate randomly for 3h. Cell trajectories were tracked and indicated. **(b)** Graph shows the mean of the track displacement from Reh cells migrating in 3D conditions as in (a). Mean n = 3 independent experiments ± SEM. **(c)** Graph shows the mean of the speed of Reh cells migrating in 3D conditions as in (a). Mean n = 3 independent experiments ± SEM. **(d)** CCRF-CEM and Reh cells pretreated or not with 1 µM OICR-9429 were fixed and actin polymerization was determined by flow cytometry. **(e)** CCRF-CEM, Reh, and primary ALL cells were lysed and subcellular localization of MLCK was resolved by western blotting. The cytosolic fraction was indicated by the loading control RhoGDI and nuclear and chromatin fractions by lamin B1. (***) P < 0.001.

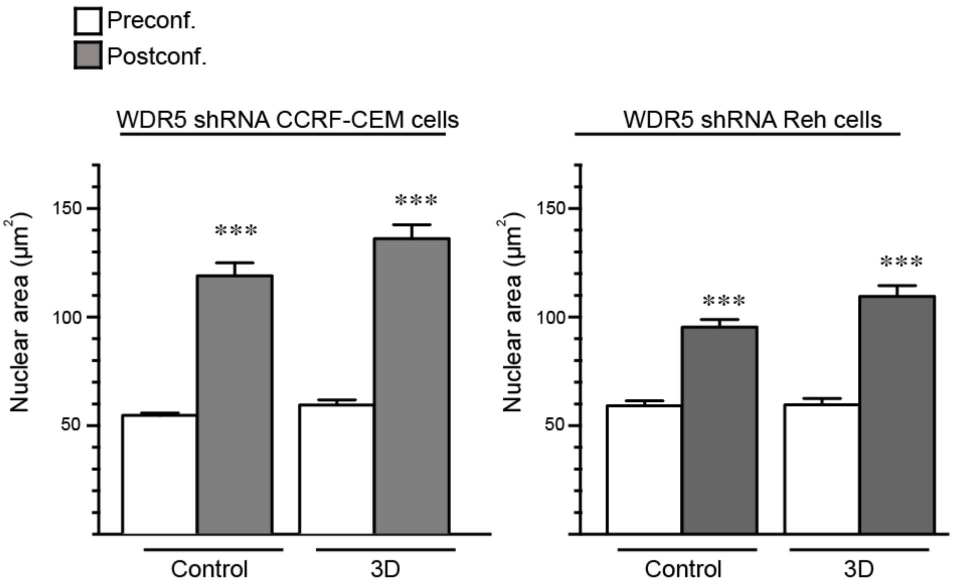


**Supplementary Fig. S6**

**Figure S6.** WDR5-depleted CCRF-CEM and Reh cells were cultured in suspension (control) or embedded in a 3D collagen matrix (3D). Then, nuclei were isolated and seeded on polylysine-coated coverslips. Confocal sections of the nuclei were taken before and after confinement. Graph shows the mean average of the nuclear area. Mean n = 15-33 nuclei ± SEM. (***) $P < 0.001$.

| Variable | Characteristics | Number of Cases | Percentage (%) |
|---|---|---|---|
| Age <10 years old | Positive | 8 | 27.6 |
| | Negative | 17 | 58.6 |
| | ND | 4 | 13.8 |
| Gender | Male | 15 | 51.7 |
| | Female | 11 | 37.9 |
| | ND | 3 | 10.3 |
| Phenotype | T-ALL | 6 | 20.7 |
| | B-ALL | 23 | 79.3 |
| High Risk Group | Positive | 6 | 20.7 |
| | Negative | 15 | 51.7 |
| | ND | 8 | 27.6 |
| WBC | Positive | 4 | 13.8 |
| | Negative | 16 | 55.2 |
| | ND | 9 | 31 |
| Blast in BM | Positive | 8 | 27.6 |
| | Negative | 4 | 13.8 |
| | ND | 17 | 58.6 |
| MRD at 15 days | Positive | 8 | 27.6 |
| | Negative | 12 | 41.4 |
| | ND | 9 | 31 |
| Relapse | Positive | 7 | 24.1 |
| | Negative | 20 | 70 |
| | ND | 2 | 6.9 |
| Death | Positive | 5 | 17.2 |
| | Negative | 19 | 65.5 |
| | ND | 5 | 17.2 |

**Table 1.** Clinical and molecular features of patients with ALL. High Risk Group is defined as poor early cytological response, or minimal residual disease level ≥ 0.05% at the end of reinduction. WBC is white blood count at diagnosis of more than 50000/µL. Blast in BM is blast in bone marrow at diagnosis of more than 80%. MRD is measurable residual disease in peripheral blood at day 15, end of induction.

| ID | 3D Avg (log2) | Control Avg (log2) | Fold Change | P-val | Gene Symbol |
|---|---|---|---|---|---|
| TC0700011571.hg.1 | 6.61 | 4.3 | 4.99 | 0.0017 | YWHAG |
| TC1000012016.hg.1 | 5.87 | 3.68 | 4.57 | 0.0014 | TIAL1 |
| TC1800008235.hg.1 | 6.38 | 4.19 | 4.56 | 0.0054 | ABHD3 |
| TC0200008536.hg.1 | 7.88 | 5.75 | 4.36 | 0.0055 | ANKRD36 |
| TC0400007460.hg.1 | 8.34 | 6.27 | 4.18 | 0.0056 | OCIAD1 |
| TC1600009183.hg.1 | 4.96 | 2.91 | 4.15 | 0.0023 | ZNF200 |
| TC2000007177.hg.1 | 6.47 | 4.51 | 3.87 | 0.0058 | RALY |
| TC0500010781.hg.1 | 5.8 | 3.89 | 3.77 | 0.0092 | SLC38A9 |
| TC0600011535.hg.1 | 7.57 | 5.67 | 3.73 | 0.0455 | RGL2 |
| TC1100011184.hg.1 | 8.9 | 7 | 3.72 | 0.0107 | SF1 |
| TC1300006811.hg.1 | 5.95 | 4.11 | 3.6 | 0.007 | PDS5B |
| TC0100008151.hg.1 | 5.83 | 4 | 3.56 | 0.0041 | RAD54L |
| TC0100015795.hg.1 | 5.35 | 3.53 | 3.53 | 0.0051 | POGZ |
| TSUnmapped00000073.hg.1 | 7.83 | 6.01 | 3.53 | 0.0068 | NDUFA10 |
| TC0800007016.hg.1 | 9.3 | 7.52 | 3.44 | 0.005 | CHMP7 |
| TC0100015791.hg.1 | 4.94 | 3.16 | 3.43 | 0.0103 | POGZ |
| TC0300010680.hg.1 | 5.14 | 3.37 | 3.41 | 0.0059 | CLASP2 |
| TC2000008237.hg.1 | 9.22 | 7.46 | 3.38 | 0.0085 | MAVS |
| TSUnmapped00000246.hg.1 | 7.62 | 5.89 | 3.3 | 0.0063 | CCDC84 |
| TC0100010926.hg.1 | 6.65 | 4.93 | 3.29 | 0.0328 | OCLM |
| TC0300007485.hg.1 | 5.32 | 3.6 | 3.29 | 0.0062 | SMIM4 |
| TC1000009922.hg.1 | 5.14 | 3.42 | 3.29 | 0.0159 | VIM |
| TC1300008080.hg.1 | 6.07 | 4.36 | 3.26 | 0.0383 | TUBGCP3 |
| TC1200009928.hg.1 | 5.14 | 3.44 | 3.25 | 0.0068 | TAS2R19 |
| TC0100015793.hg.1 | 5.2 | 3.5 | 3.24 | 0.0061 | POGZ |
| TSUnmapped00000538.hg.1 | 6.58 | 4.9 | 3.2 | 0.0286 | DGKD |
| TC1700008441.hg.1 | 6.13 | 4.45 | 3.2 | 0.0084 | PPM1E |
| TC0200014738.hg.1 | 5.44 | 3.78 | 3.15 | 0.0076 | RBMS1 |
| TC1600007741.hg.1 | 4.59 | 2.95 | 3.13 | 0.007 | PHKB |
| TSUnmapped00000262.hg.1 | 6.41 | 4.78 | 3.1 | 0.0313 | MLXIP |
| TC1000010918.hg.1 | 5.02 | 3.4 | 3.09 | 0.0079 | LRRC20 |
| TC0900006539.hg.1 | 5.81 | 4.19 | 3.07 | 0.018 | JAK2 |
| TSUnmapped00000725.hg.1 | 5.01 | 3.39 | 3.07 | 0.0154 | CCDC84 |
| TC0100012695.hg.1 | 5.05 | 3.45 | 3.04 | 0.021 | DNAJC11 |
| TC1200011313.hg.1 | 6.59 | 5 | 3.02 | 0.0139 | NAP1L1 |
| TC1000007885.hg.1 | 5.96 | 4.36 | 3.02 | 0.0084 | SRGN |
| TC0300013636.hg.1 | 4.54 | 2.97 | 2.98 | 0.0168 | ACAP2 |
| TC1200010686.hg.1 | 6.33 | 4.76 | 2.97 | 0.0331 | SLC11A2 |
| TC0500009423.hg.1 | 6.42 | 4.85 | 2.97 | 0.0088 | NPM1 |
| TC0300007014.hg.1 | 6.11 | 4.56 | 2.93 | 0.015 | ARPP21 |
| TC0X00008383.hg.1 | 5.02 | 3.47 | 2.92 | 0.0122 | OCRL |
| TSUnmapped00000041.hg.1 | 4.84 | 3.3 | 2.9 | 0.0204 | ATG16L1 |
| TC1300008706.hg.1 | 5.64 | 4.12 | 2.87 | 0.0388 | ELF1 |
| TC0900007100.hg.1 | 4.99 | 3.47 | 2.86 | 0.0375 | NPR2 |
| TC0200010729.hg.1 | 7.04 | 5.53 | 2.85 | 0.0222 | XRCC5 |
| TC0100013889.hg.1 | 4.82 | 3.32 | 2.83 | 0.041 | ZMYND12 |
| TC0300007016.hg.1 | 6.35 | 4.85 | 2.82 | 0.016 | ARPP21 |
| TC0100014286.hg.1 | 4.71 | 3.21 | 2.81 | 0.0388 | USP24 |
| TC0100015947.hg.1 | 12.04 | 10.56 | 2.8 | 0.0145 | THBS3 |
| TC0200013030.hg.1 | 5.48 | 4 | 2.79 | 0.0391 | DYSF |

| | | | | | |
|---|---|---|---|---|---|
| TC2000009886.hg.1 | 6.76 | 5.29 | 2.78 | 0.0274 | PANK2 |
| TC0100011219.hg.1 | 6.21 | 4.74 | 2.77 | 0.0387 | PPP1R12B |
| TC0400009766.hg.1 | 5.85 | 4.37 | 2.77 | 0.0411 | MXD4 |
| TC1300008705.hg.1 | 5.08 | 3.61 | 2.77 | 0.0125 | ELF1 |
| TC1600011365.hg.1 | 9.98 | 8.52 | 2.76 | 0.0151 | NPIPB9 |
| TC0100009149.hg.1 | 7.18 | 5.71 | 2.76 | 0.0336 | PTBP2 |
| TC0200008534.hg.1 | 9.65 | 8.2 | 2.74 | 0.0149 | ANKRD36 |
| TC0100014459.hg.1 | 4.74 | 3.29 | 2.74 | 0.0195 | LINC01359 |
| TC0500013064.hg.1 | 5.43 | 3.98 | 2.73 | 0.0359 | MGAT4B |
| TC1500007805.hg.1 | 5.16 | 3.72 | 2.72 | 0.0285 | BBS4 |
| TC0900009679.hg.1 | 4.03 | 2.59 | 2.72 | 0.0299 | MLLT3 |
| TSUnmapped00000692.hg.1 | 4.47 | 3.03 | 2.7 | 0.0449 | ZNF502 |
| TC0100008130.hg.1 | 5.17 | 3.74 | 2.7 | 0.0207 | NASP |
| TC1900011796.hg.1 | 6.06 | 4.63 | 2.69 | 0.0215 | NDUFA3 |
| TC0900008628.hg.1 | 6.8 | 5.37 | 2.68 | 0.0398 | CNTRL |
| TC0100012335.hg.1 | 4.81 | 3.39 | 2.68 | 0.0151 | OR2AK2 |
| TC0500009012.hg.1 | 5.79 | 4.37 | 2.67 | 0.0189 | TCERG1 |
| TC0700013583.hg.1 | 6.5 | 5.08 | 2.67 | 0.018 | MTERF1 |
| TC0600014111.hg.1 | 9.55 | 8.13 | 2.66 | 0.0372 | SYNGAP1 |
| TC1700007897.hg.1 | 6.97 | 5.56 | 2.66 | 0.0289 | HSD17B1 |
| TC1600011505.hg.1 | 10.89 | 9.49 | 2.64 | 0.0383 | NPIPB4 |
| TC0X00008138.hg.1 | 5.01 | 3.61 | 2.64 | 0.0197 | ALG13 |
| TC1700011495.hg.1 | 5.26 | 3.88 | 2.61 | 0.0481 | APOH |
| TC1000007510.hg.1 | 9.03 | 7.65 | 2.6 | 0.0177 | AGAP9 |
| TC0400009684.hg.1 | 6.41 | 5.04 | 2.6 | 0.0316 | ZNF721 |
| TC1600008407.hg.1 | 9.92 | 8.55 | 2.59 | 0.0185 | NPIPB15 |
| TC0500009103.hg.1 | 4.72 | 3.34 | 2.59 | 0.02 | IRGM |
| TC0100008026.hg.1 | 7.91 | 6.53 | 2.59 | 0.0221 | YBX1 |
| TC0600013180.hg.1 | 4.83 | 3.46 | 2.58 | 0.0284 | CTAGE9 |
| TC1200007421.hg.1 | 6.02 | 4.66 | 2.58 | 0.0183 | ANO6 |
| TC0200009865.hg.1 | 5.07 | 3.71 | 2.57 | 0.0238 | SCN2A |
| TSUnmapped00000053.hg.1 | 7.01 | 5.65 | 2.57 | 0.0286 | RPL7A |
| TC1700012027.hg.1 | 7.27 | 5.91 | 2.57 | 0.0313 | ENTHD2 |
| TC1900011328.hg.1 | 11.66 | 10.31 | 2.55 | 0.0192 | ZNF160 |
| TC1900011913.hg.1 | 5.8 | 4.45 | 2.55 | 0.0325 | LINC00663 |
| TC0800006884.hg.1 | 6.11 | 4.76 | 2.54 | 0.0315 | PCM1 |
| TC0200014736.hg.1 | 5.4 | 4.05 | 2.54 | 0.0362 | RBMS1 |
| TC0700007201.hg.1 | 5.19 | 3.85 | 2.54 | 0.0359 | ANLN |
| TC1000010509.hg.1 | 9.6 | 8.27 | 2.53 | 0.0203 | AGAP4 |
| TC0100008129.hg.1 | 4.72 | 3.39 | 2.52 | 0.0304 | NASP |
| TC0800011595.hg.1 | 5.62 | 4.28 | 2.52 | 0.0201 | EXT1 |
| TC2000008940.hg.1 | 6.51 | 5.18 | 2.52 | 0.0469 | PIGU |
| TC1500010947.hg.1 | 7.11 | 5.78 | 2.52 | 0.0221 | TM2D3 |
| TC1600007353.hg.1 | 11.13 | 9.8 | 2.51 | 0.0281 | NPIPB8 |
| TC2200007493.hg.1 | 12.28 | 10.96 | 2.49 | 0.0244 | MEI1 |
| TC1400010170.hg.1 | 5.4 | 4.08 | 2.49 | 0.0295 | SETD3 |
| TC1200009183.hg.1 | 7.98 | 6.66 | 2.48 | 0.0496 | SETD1B |
| TC0800007767.hg.1 | 4.89 | 3.58 | 2.48 | 0.0275 | RAB2A |
| TC0100007800.hg.1 | 6.17 | 4.87 | 2.47 | 0.0298 | THRAP3 |
| TC0X00006625.hg.1 | 3.97 | 2.66 | 2.47 | 0.0409 | TLR7 |
| TC1300009714.hg.1 | 9.25 | 7.95 | 2.46 | 0.0312 | ARGLU1 |

| | | | | | |
|---|---|---|---|---|---|
| TSUnmapped00000212.hg.1 | 6.16 | 4.86 | 2.46 | 0.0399 | MLXIP |
| TC1800006679.hg.1 | 8.04 | 6.75 | 2.45 | 0.0287 | NAPG |
| TC1000007598.hg.1 | 9.75 | 8.47 | 2.44 | 0.0242 | AGAP6 |
| TC1100012000.hg.1 | 5.24 | 3.95 | 2.44 | 0.0441 | GPR83 |
| TC1600011351.hg.1 | 7.28 | 6 | 2.43 | 0.0447 | CARHSP1 |
| TC0300014054.hg.1 | 4.77 | 3.49 | 2.43 | 0.025 | PIK3CB |
| TC0200016582.hg.1 | 6.99 | 5.71 | 2.42 | 0.0302 | NABP1 |
| TC0100015796.hg.1 | 6.4 | 5.12 | 2.42 | 0.0322 | POGZ |
| TC1900008925.hg.1 | 6.28 | 5.01 | 2.41 | 0.0472 | U2AF2 |
| TC0500006648.hg.1 | 9.45 | 8.18 | 2.41 | 0.0314 | PAPD7 |
| TSUnmapped00000085.hg.1 | 5.28 | 4.02 | 2.41 | 0.0274 | CCDC84 |
| TC1200010752.hg.1 | 4.71 | 3.44 | 2.41 | 0.0383 | KRT84 |
| TC1700010700.hg.1 | 7.2 | 5.94 | 2.4 | 0.0419 | DNAJC7 |
| TC1800006513.hg.1 | 6.81 | 5.55 | 2.4 | 0.0328 | TGIF1 |
| TC1200008686.hg.1 | 7.75 | 6.49 | 2.39 | 0.0497 | CHST11 |
| TC0100017436.hg.1 | 5.44 | 4.18 | 2.39 | 0.0375 | TAF1A |
| TC0100015771.hg.1 | 6.44 | 5.19 | 2.39 | 0.028 | SEMA6C |
| TC0200016739.hg.1 | 4.46 | 3.21 | 2.38 | 0.0275 | ARL5A |
| TSUnmapped00000222.hg.1 | 6.07 | 4.82 | 2.38 | 0.0308 | ZNF780A |
| TC1700010703.hg.1 | 6.74 | 5.49 | 2.38 | 0.0443 | NKIRAS2 |
| TSUnmapped00000149.hg.1 | 4.45 | 3.21 | 2.37 | 0.0498 | ZNF35 |
| TC0200009774.hg.1 | 6.06 | 4.82 | 2.37 | 0.029 | TANC1 |
| TC0100007305.hg.1 | 5.08 | 3.84 | 2.37 | 0.0281 | KDM1A |
| TC0300013601.hg.1 | 4.48 | 3.24 | 2.37 | 0.0401 | ATP13A3 |
| TC1500008346.hg.1 | 5.72 | 4.49 | 2.36 | 0.0415 | PRC1-AS1 |
| TC2200009128.hg.1 | 7.57 | 6.34 | 2.36 | 0.0365 | ALG12 |
| TC1200006688.hg.1 | 4.63 | 3.4 | 2.34 | 0.0499 | NANOG |
| TC1000011660.hg.1 | 5.45 | 4.23 | 2.34 | 0.0335 | MGEA5 |
| TC0400012916.hg.1 | 5 | 3.78 | 2.33 | 0.0471 | ARAP2 |
| TC1400010759.hg.1 | 5.02 | 3.8 | 2.32 | 0.0352 | ATP6V1D |
| TC0600011723.hg.1 | 4.08 | 2.87 | 2.31 | 0.0388 | BTBD9 |
| TC1000009420.hg.1 | 5.1 | 3.91 | 2.29 | 0.0441 | INPP5A |
| TC1600007037.hg.1 | 10.81 | 9.62 | 2.29 | 0.0374 | NPIPA7 |
| TC0X00008514.hg.1 | 6.54 | 5.35 | 2.28 | 0.0373 | DDX26B |
| TC0100015797.hg.1 | 5.72 | 4.53 | 2.28 | 0.0375 | POGZ |
| TC0100018298.hg.1 | 8.66 | 7.48 | 2.27 | 0.0493 | INTS3 |
| TC1200006647.hg.1 | 7.44 | 6.26 | 2.27 | 0.0389 | GNB3 |
| TC0300009224.hg.1 | 5.07 | 3.9 | 2.26 | 0.0495 | P2RY1 |
| TC0800006697.hg.1 | 5.1 | 3.93 | 2.26 | 0.0483 | MSRA |
| TC0100016356.hg.1 | 4.45 | 3.27 | 2.26 | 0.0452 | F5 |
| TC0400009198.hg.1 | 5.31 | 4.14 | 2.25 | 0.0492 | APELA |
| TC0300007165.hg.1 | 10.95 | 9.78 | 2.25 | 0.0441 | NKTR |
| TC1100009859.hg.1 | 5.83 | 4.66 | 2.25 | 0.0423 | NUP98 |
| TC1200012593.hg.1 | 5.97 | 4.8 | 2.24 | 0.0385 | CLEC4A |
| TSUnmapped00000239.hg.1 | 4.74 | 3.58 | 2.24 | 0.0415 | ATG16L1 |
| TC1100013006.hg.1 | 5.21 | 4.05 | 2.24 | 0.0387 | CNTF |
| TC0800011566.hg.1 | 6.12 | 4.96 | 2.23 | 0.0424 | RAD21 |
| TC1000012573.hg.1 | 6.03 | 4.87 | 2.23 | 0.037 | MMRN2 |
| TSUnmapped00000670.hg.1 | 6.07 | 4.91 | 2.23 | 0.0424 | ZNF780A |
| TC1000006907.hg.1 | 8.59 | 7.44 | 2.21 | 0.0396 | STAM |
| TC0800012076.hg.1 | 7.31 | 6.17 | 2.21 | 0.0473 | ARC |

| | | | | | |
|---|---|---|---|---|---|
| TC1300008983.hg.1 | 7.34 | 6.2 | 2.2 | 0.0444 | INTS6 |
| TC0100008772.hg.1 | 7.26 | 6.12 | 2.2 | 0.042 | RABGGTB |
| TC2000008299.hg.1 | 7.14 | 6.01 | 2.19 | 0.0418 | GPCPD1 |
| TC1100013035.hg.1 | 4.8 | 3.67 | 2.19 | 0.0499 | PPP2R5B |
| TC0200011014.hg.1 | 7.66 | 6.54 | 2.18 | 0.0497 | FBXO36 |
| TC1400006691.hg.1 | 6.93 | 5.81 | 2.18 | 0.0416 | THTPA |
| TC0800009878.hg.1 | 8.81 | 7.7 | 2.17 | 0.0456 | ENTPD4 |
| TC1900008827.hg.1 | 6.65 | 5.53 | 2.16 | 0.0436 | CACNG8 |
| TC1200012827.hg.1 | 4.31 | 3.21 | 2.15 | 0.0474 | CCER1 |
| TC0700009400.hg.1 | 5.27 | 4.17 | 2.14 | 0.0445 | TAS2R5 |
| TC1600009621.hg.1 | 10.84 | 9.76 | 2.12 | 0.0491 | SMG1 |
| TC0300013820.hg.1 | 4.55 | 3.47 | 2.12 | 0.0481 | ZNF660 |
| TC1900008015.hg.1 | 4.68 | 3.61 | 2.11 | 0.0491 | SPRED3 |
| TC0X00010406.hg.1 | 5.99 | 7.08 | -2.14 | 0.0485 | TCEAL8 |
| TC0X00011309.hg.1 | 7.12 | 8.25 | -2.18 | 0.05 | HNRNPH2 |
| TC1000007986.hg.1 | 7.59 | 8.75 | -2.24 | 0.0471 | ANAPC16 |
| TSUnmapped00000798.hg.1 | 4.11 | 5.28 | -2.25 | 0.0404 | HMBS |
| TC1700007153.hg.1 | 5.36 | 6.54 | -2.26 | 0.0344 | GRAPL |
| TC0500007935.hg.1 | 6.39 | 7.57 | -2.26 | 0.0493 | ZCCHC9 |
| TC1600011573.hg.1 | 6.05 | 7.25 | -2.3 | 0.0435 | GCSH |
| TC0100012828.hg.1 | 6.18 | 7.39 | -2.31 | 0.0489 | LZIC |
| TC1100007168.hg.1 | 5.79 | 7.01 | -2.33 | 0.0425 | ARL14EP |
| TC0X00008097.hg.1 | 6.25 | 7.47 | -2.33 | 0.0413 | NXT2 |
| TC1400009065.hg.1 | 6.12 | 7.35 | -2.33 | 0.0425 | FKBP3 |
| TC1700009551.hg.1 | 4.79 | 6.02 | -2.34 | 0.0429 | ZNF232 |
| TC2100008499.hg.1 | 7.32 | 8.55 | -2.36 | 0.0365 | SMIM11A |
| TC2000007117.hg.1 | 8.62 | 9.88 | -2.39 | 0.0481 | ASXL1 |
| TC2000006580.hg.1 | 4.59 | 5.85 | -2.4 | 0.0431 | RASSF2 |
| TC0700012011.hg.1 | 4.15 | 5.41 | -2.4 | 0.0303 | ACTL6B |
| TC2200008637.hg.1 | 4.6 | 5.87 | -2.41 | 0.0292 | IL2RB |
| TC1400010730.hg.1 | 4.11 | 5.39 | -2.43 | 0.0253 | EGLN3 |
| TC0200016703.hg.1 | 6.78 | 8.06 | -2.44 | 0.0347 | CHMP3 |
| TC1400009104.hg.1 | 8.83 | 10.12 | -2.44 | 0.0396 | RPL36AL |
| TC2100006460.hg.1 | 7.26 | 8.57 | -2.48 | 0.0326 | SMIM11A |
| TC0200016584.hg.1 | 7.22 | 8.53 | -2.48 | 0.0425 | HSPE1 |
| TC1400006715.hg.1 | 5.28 | 6.59 | -2.48 | 0.0438 | PCK2 |
| TC0600008209.hg.1 | 5.78 | 7.11 | -2.5 | 0.0468 | CENPQ |
| TC0800011239.hg.1 | 7.95 | 9.28 | -2.52 | 0.0356 | COX6C |
| TC1500010867.hg.1 | 6.48 | 7.82 | -2.53 | 0.0301 | PIGBOS1 |
| TC0200016455.hg.1 | 5.71 | 7.05 | -2.54 | 0.0438 | C2orf74 |
| TC0900006969.hg.1 | 7.07 | 8.43 | -2.56 | 0.0459 | CHMP5 |
| TC0700009676.hg.1 | 7.38 | 8.74 | -2.58 | 0.0392 | GIMAP7 |
| TC0200006788.hg.1 | 5.29 | 6.67 | -2.6 | 0.0338 | DDX1 |
| TC2200006654.hg.1 | 5.41 | 6.8 | -2.62 | 0.019 | ZNF74 |
| TC1300006729.hg.1 | 7.18 | 8.58 | -2.64 | 0.0473 | POMP |
| TC1500009438.hg.1 | 4.57 | 5.97 | -2.64 | 0.0326 | LYSMD2 |
| TC1000007883.hg.1 | 6.86 | 8.26 | -2.64 | 0.0341 | SRGN |
| TC1100008991.hg.1 | 5 | 6.4 | -2.65 | 0.0399 | DDX10 |
| TC0200009813.hg.1 | 7.62 | 9.04 | -2.68 | 0.0383 | PSMD14 |
| TC1700010551.hg.1 | 9.94 | 11.38 | -2.71 | 0.0372 | RPL23 |
| TC0200014870.hg.1 | 7.66 | 9.11 | -2.72 | 0.0358 | METTL5 |

| | | | | | |
|---|---|---|---|---|---|
| TC1300006699.hg.1 | 6.34 | 7.79 | -2.74 | 0.0263 | ATP5EP2 |
| TC0100011533.hg.1 | 6.84 | 8.29 | -2.74 | 0.0412 | ATF3 |
| TC1200010454.hg.1 | 8.33 | 9.79 | -2.75 | 0.0432 | ZCRB1 |
| TC1700008282.hg.1 | 7.62 | 9.09 | -2.76 | 0.0332 | NME1 |
| TC1500007056.hg.1 | 5.9 | 7.37 | -2.77 | 0.0182 | ADAL |
| TC0100011012.hg.1 | 5.52 | 6.99 | -2.78 | 0.0178 | RGS1 |
| TC0X00011310.hg.1 | 8.51 | 9.99 | -2.8 | 0.0262 | RPL36A-HNRNPH2 |
| TC0800008120.hg.1 | 4.13 | 5.62 | -2.8 | 0.0473 | LRRCC1 |
| TC0300014034.hg.1 | 7.53 | 9.02 | -2.81 | 0.0257 | COX17 |
| TC1000010452.hg.1 | 5 | 6.52 | -2.87 | 0.0365 | ZNF32 |
| TC0900011428.hg.1 | 6.07 | 7.59 | -2.87 | 0.0291 | NDUFA8 |
| TC2000006867.hg.1 | 5.08 | 6.61 | -2.88 | 0.0277 | NAA20 |
| TC1600011416.hg.1 | 5.12 | 6.66 | -2.92 | 0.0359 | DUS2 |
| TC0500012070.hg.1 | 6.95 | 8.5 | -2.93 | 0.0205 | CDKN2AIPNL |
| TC0500010717.hg.1 | 6.81 | 8.37 | -2.96 | 0.0496 | MOCS2 |
| TC0300013146.hg.1 | 7.42 | 8.99 | -2.98 | 0.0296 | TNFSF10 |
| TC1200008703.hg.1 | 5.38 | 6.95 | -2.99 | 0.0305 | C12orf45 |
| TC0100014014.hg.1 | 8.5 | 10.08 | -2.99 | 0.0434 | PRDX1 |
| TC0500007966.hg.1 | 6.44 | 8.03 | -3 | 0.0422 | XRCC4 |
| TC0500007667.hg.1 | 6.39 | 7.98 | -3.02 | 0.0252 | MRPS36 |
| TC1500010893.hg.1 | 5.64 | 7.23 | -3.02 | 0.0401 | PTPN9 |
| TC0600007203.hg.1 | 6.58 | 8.18 | -3.04 | 0.0353 | ACOT13 |
| TC0500010920.hg.1 | 8.03 | 9.67 | -3.12 | 0.0231 | SREK1IP1 |
| TC0300013960.hg.1 | 6.98 | 8.62 | -3.13 | 0.0328 | HIGD1A |
| TC0400012930.hg.1 | 6.03 | 7.7 | -3.17 | 0.0337 | COX18 |
| TC1000011716.hg.1 | 7.08 | 8.75 | -3.19 | 0.0482 | USMG5 |
| TC0600014068.hg.1 | 7.22 | 8.9 | -3.2 | 0.0485 | TMEM14C |
| TC0700010247.hg.1 | 5.9 | 7.58 | -3.2 | 0.042 | RPA3 |
| TC0100008152.hg.1 | 6.78 | 8.49 | -3.28 | 0.0406 | UQCRH |
| TC0200006440.hg.1 | 7.16 | 8.93 | -3.4 | 0.0234 | ACP1 |
| TC0600014292.hg.1 | 6.56 | 8.32 | -3.4 | 0.0192 | OARD1 |
| TC0100012020.hg.1 | 5.94 | 7.73 | -3.45 | 0.0362 | COA6 |
| TC1300008439.hg.1 | 6.24 | 8.07 | -3.55 | 0.0415 | MTIF3 |
| TC0300013795.hg.1 | 6.46 | 8.29 | -3.56 | 0.0369 | OXNAD1 |
| TC0X00010172.hg.1 | 6.44 | 8.27 | -3.56 | 0.0067 | HMGN5 |
| TC0600009142.hg.1 | 6.96 | 8.9 | -3.83 | 0.0289 | RPF2 |
| TC1500009738.hg.1 | 7.13 | 9.15 | -4.05 | 0.0324 | KIAA0101 |
| TC1300009273.hg.1 | 8.21 | 10.26 | -4.17 | 0.0392 | COMMD6 |
| TC1200008591.hg.1 | 6.57 | 8.64 | -4.2 | 0.0133 | ACTR6 |
| TC0600007263.hg.1 | 6.41 | 8.5 | -4.26 | 0.0193 | HIST1H4A |
| TC0200010438.hg.1 | 6.28 | 8.38 | -4.31 | 0.0339 | NDUFB3 |
| TC2000007792.hg.1 | 6.59 | 8.72 | -4.38 | 0.0182 | PFDN4 |
| TC1100009040.hg.1 | 6.54 | 8.88 | -5.06 | 0.0481 | C11orf1 |
| TC1100013147.hg.1 | 9.17 | 11.56 | -5.26 | 0.0349 | RPS13 |
| TC0200013095.hg.1 | 5.63 | 8.28 | -6.29 | 0.023 | BOLA3 |
| TC2200009235.hg.1 | 6.47 | 9.22 | -6.72 | 0.011 | SNRPD3 |
